# The Kink-Turn Motif: A Powerful Test for Revealing Weaknesses in RNA Force Fields

**DOI:** 10.1101/2025.05.13.653794

**Authors:** Toon Lemmens, Vojtěch Mlýnský, Jiří Šponer, Martin Pykal, Pavel Banáš, Michal Otyepka, Miroslav Krepl

**Author notes:** Corresponding author: Miroslav Krepl.

## Abstract

The kink-turn is a recurrent RNA structural motif that induces a sharp bend (kink) in the A-form RNA helix. It is defined by key structural features, including consecutive sheared AG base pairs, an A-minor interaction, and multiple base-sugar interactions. Accurate representation of these densely packed non-canonical motifs in molecular dynamics simulations poses a significant challenge for contemporary force fields (FFs). Here, we present extended simulations of ribosomal kink-turn 7 (Kt-7) using a broad spectrum of pair-additive and polarizable RNA FFs. None of the tested FFs manage to flawlessly describe all the specific structural features of the Kt-7, which are described in detail in this work. Still, several FFs provide rather acceptable results and should not cause problems in simulations of larger RNAs containing a kink-turn. On aggregate, the widely used OL3 (ff99bsc0χ_OL3_) and polarizable AMOEBA FFs achieve the best performance. On the other hand, some more recently parametrized FF variants struggle to describe the Kt-7’s tertiary A-minor interaction – an ubiquitous tertiary contact in RNA. This raises some concerns about the broader applicability of these FFs and suggests that they may be overfitted to small RNA model systems, such as RNA tetranucleotides. The difficulties manifest as either reversible local disruptions of the A-minor interaction and other tertiary contacts or, in some cases, irreversible unkinking of the entire motif. Based on our findings, we strongly suggest including the kink-turn motif in training and benchmarking datasets as the quintessential regression test to enhance the robustness and accuracy of RNA FF parametrization efforts.

## Introduction

The kink-turn is a recurrent structural motif, widespread in RNA across all domains of life,^1, 2^ where it contributes to a myriad of structural and biological functions. Kink-turns mediate protein-RNA and RNA-RNA tertiary contacts in ribosomes,^3^ form crucial structural element in several riboswitches,^4–6^ and are involved in the assembly of the eukaryotic spliceosome.^7, 8^ They are a powerful tool in RNA nanotechnology, allowing for precise control over RNA folding and modular assembly in RNA origami and nanostructures.^9^ While their characteristic bent shape is strongly conserved, the individual kink-turns display large sequence variations, especially in the bulge region.^10^ Closest to the “consensual” sequence is the kink-turn 7 from the large ribosomal subunit of *Haloarcula marismortui* (Figure 1; henceforth referred to as Kt-7),^1^ making it the most extensively studied kink-turn. Kt-7 also plays a role in the assembly of the box C/D and H/ACA snoRNP complexes, as well as methylation and pseudouridylation of archaeal and eukaryotic RNAs.^11–14^ The Kt-7 structure is formed by a longer (b) and shorter (n) RNA strand, with the longer strand containing the three-nucleotide GAA bulge (loop). The structure further includes the canonical (C) and non-canonical (NC) stems separated by the bulge and containing the characteristic cis-Watson-Crick (*c*WW, canonical) GC and trans-Hoogsteen-sugar edge (*t*HS) AG base pairs, respectively.^15^ The bent shape of the kink-turn is stabilized by essential tertiary interactions formed among the two stems and the bulge (Figure 1).^3^

**Figure 1:**
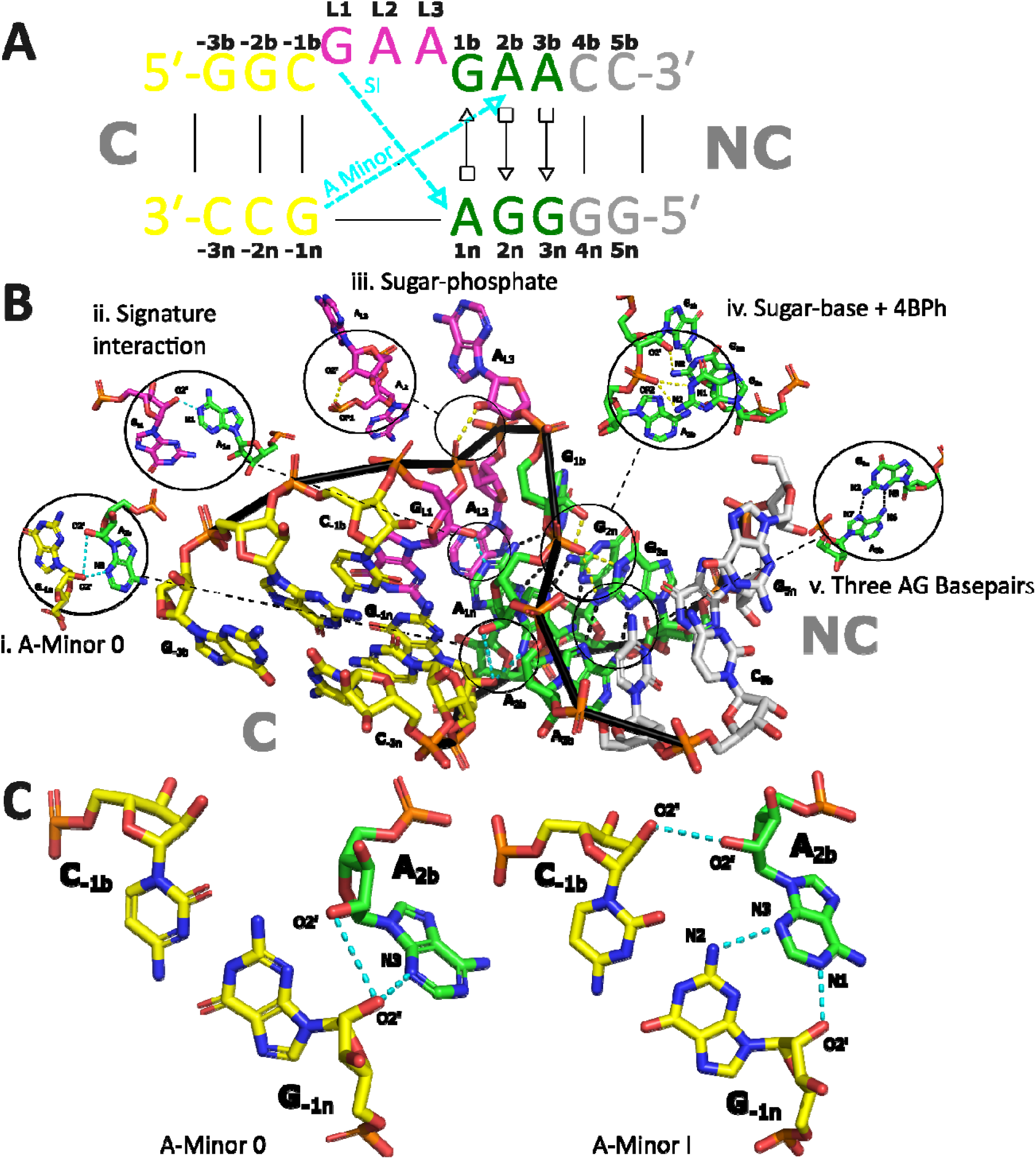
Structural overview of the Kink-turn 7 (Kt-7) from *Haloarcula marismortui*. (A) Schematic representation of Kt-7 using the Leontis & Westhof base pair classification.^15^ The canonical stem (*C*), the AG base pairs, the Watson-Crick base pairs of the non-canonical stem (*NC*) and the bulge are in yellow, green, gray and purple, respectively. The same color coding will be used throughout this work. The main tertiary interactions stabilizing the kink-turn – signature interaction (SI) and A-minor are in cyan. The nucleotides are labeled according to their established annotation^1^ which will be used throughout the study. (B) The truncated X-ray structure of isolated Kt-7 (PDB: 4C40)^19^ with carbon-atom color-coding as in A. The H-bonds are indicated by dashed lines. The base-phosphate type 4 (4BPh)^20^, sugar-phosphate and sugar-base interactions are in yellow, and the AG base pair H-bonds are in black. The RNA backbone is highlighted with a black ribbon. The inset Figures i-v show selected interactions in greater detail. (C) Comparison between the A-minor 0 and A-minor I interaction.

The first of these essential interactions is known as the kink-turn’s signature interaction (SI). It is formed between atoms G_L1_(O2’) and A_1n_(N1), and is utterly indispensable for the folding of all kink-turns (Figure 1B).^3^ Secondly, there is an A-minor interaction between A_2b_ and the G_-1n_:C_-1b_ base pair.^3^ The A-minor interaction, where adenine is inserted into a minor groove of canonical A-RNA duplex, is not only a critical element for the kink-turn motifs but also the most widespread tertiary interaction in RNA.^16, 17^ The A-minor interaction can adopt four distinct substates (A-minor type 0, I, II, and III) that differ by their spatial orientation and H-bonding patterns.^18^ Generally, only types 0 and I A-minor interactions are observed in kink-turns (Figure 1C). For Kt-7, the type I is experimentally observed when the kink-turn is embedded in its ribosomal context while the type 0 is observed for isolated Kt-7 or when complexed with the L7Ae protein.^19^ In addition to the SI and A-minor interactions, there are additional tertiary contacts stabilizing Kt-7. Namely the 4BPh base-phosphate interaction^20^ between G_3n_ and A_2b_, the sugar-phosphate interaction between A_L2_ and A_L3_, and the sugar-base interaction between G_1b_ and G_2n_ (Figure 1B). Kt-7 also adopts characteristic, non-canonical backbone conformations, with several backbone dihedral suites exhibiting conserved non-canonical substates (Supporting Information Table S1).^21, 22^

Molecular dynamics (MD) simulations are a computational method for observing the thermal fluctuations (dynamics) of biomolecular systems with an essentially unlimited spatiotemporal resolution. The atomic movements are described using carefully calibrated empirical potentials, collectively referred to as the force fields (FFs).^23^ Assuming a high-quality starting structure, the success or failure of the MD simulations to reproduce real biomolecular dynamics rests on the quality of the utilized FF. In case of RNA, a persistent challenge lies in developing a FF capable of accurately describing both single-stranded and structured RNAs.^24^ Despite over a decade of dedicated efforts,^25–43^ this remains an open-ended question, with uncertainty as to whether such a comprehensive solution is even achievable.^44^ As occasionally noted earlier, the Kt-7 could be a useful benchmarking system for RNA FF testing.^45–47^ Its densely packed network of non-canonical interactions makes it highly sensitive to even relatively small FF imbalances, especially those arising from efforts to stabilize the single-stranded A-RNA conformations which have been common target structures in recent RNA FF development attempts.^26, 29, 48–50^ This could make Kt-7 a potentially invaluable model for regression testing to detect potential FF overfitting.

In this work, we perform extensive benchmarking of series of modern RNA FFs, namely the pairwise additive OL3 (ff99bsc0χ_OL3_),^51–53^ DESRES,^54^ DES-Amber,^47^ ROC,^55^ PAK,^56^ BSSF1,^57^ Chen&Garcia,^58^ CHARMM36,^59^ and the polarizable FFs CHARMM_Drude_^60–63^ and AMOEBA.^64, 65^ We have further tested two extensions of the OL3 FF abbreviated as OL3_0BPh,CP_-gHBfix21 and OL3_R2.7_, which are further described in the Methods section. Our study unveils a very diverse performance with some FFs providing entirely stable trajectories and others leading to loss of the kinked shape (straightening) of the kink-turn. However, to some extent, all tested FFs struggled with one or more Kt-7 interactions, even though most preserved its kinked shape. Taken together, our data highlight Kt-7 as a highly advantageous model for evaluating the balance of interactions in RNA FFs. The rich equilibrium dynamics of Kt-7, including transitions between the two types of A-minor interactions,^66, 67^ means that many types of FF imbalances lead to relatively swift occurrence of structural problems, well within the timescales easily accessible by standard MD simulations on contemporary GPU-accelerated hardware. This eliminates the need for enhanced sampling simulations when benchmarking Kt-7. In addition, the Kt-7 autonomously folds in the presence of monovalent ions^2, 68^ and does not require Mg^2+^, whose computational modeling can pose further challenges.^24^ In this study we identify the key structural features that should be monitored during FF benchmarking on Kt-7. We also suggest that it should serve as an indispensable model for FF testing due to the many potential pitfalls in simulations and its distinctiveness from typical training sets, which primarily consist of tetranucleotides and tetraloops.^27, 29^ Any new RNA FF claiming general applicability should demonstrate performance on Kt-7 that matches or exceeds previous parametrizations, to avoid imbalance or overfitting to a specific structural motif or interaction.

## Materials and methods

### Starting structure selection and RNA FFs

We used the X-ray structure of isolated Kt-7 (PDB: 4C40)^19^ as the starting structure for almost all our simulations. A single kink-turn molecule was extracted from the asymmetric unit and truncated as shown in Figure 1A. The tested RNA FFs included non-polarizable FFs OL3,^51–53^ DESRES,^54^ DES-Amber,^47^ ROC,^55^ PAK,^56^ BSSF1,^57^ Chen&Garcia^58^ and CHARMM36,^59^ as well as polarizable FFs CHARMM ^60–63^ and AMOEBA.^64, 65^ We additionally tested two recently proposed modifications of the non-bonded terms in the standard OL3 FF, abbreviated as OL3_0BPh,CP_-gHBfix21 and OL3_R2.7_. The OL3_0BPh,CP_-gHBfix21 variant primarily incorporates the gHBfix21 potential, which was developed using a machine-learning approach trained on experimental data for RNA tetranucleotides and tetraloops.^41^ This potential fine-tunes H-bond interactions and was parameterized together with the OPC water model and the so called Case phosphates (CP) modification, a refinement of phosphate oxygen Lennard-Jones (LJ) parameters introduced by Steinbrecher et al.^69^ The CP modification was applied for the OL3_0BPh,CP_-gHBfix21 in the present study, alongside our correction of the α, γ, δ, and ζ backbone dihedral potentials, which compensates for the altered phosphate LJ parameters.^70^ Additionally, we incorporated an independently developed non-bonded fix (NBfix) correction, NBfix_0BPh_, which adjusts the LJ potential for -H8…O5’ and - H6…O5’ atom pairs.^43^ This correction eliminates steric clashes in intranucleotide type 0 base-phosphate (0BPh) interactions. The OL3_R2.7_ variant applies a similar NBfix correction to almost all C-H…O interactions in RNA, universally reducing the H…O R_min_ LJ distance to 2.7 Å.^71^ This “hydrogen repulsion modification” was originally designed to mitigate C-H…O steric conflicts in RNA hairpin loops but it also introduces spurious C-H…O interactions in other regions of RNA structures.^72^ Lastly, we have also tested the standard OL3 FF on the Kt-7 structure extracted from the *Haloarcula marismortui* large ribosomal subunit (residues 76-83 and 91-101; PDB: 1S72),^73^ which has the A-minor I starting conformation instead of the A-minor 0 (see Introduction). These simulations are evaluated separately from the rest, and we are not referring to them unless explicitly stated.

### Water models and ion parameters

When selecting the water model and ion parameters, we primarily followed the specific recommendations in the original literature made by FF authors. When none were given, we made a choice reflecting the typical use patterns observed in various studies. To that end, we used OPC^74^ with OL3, OL3_0BPh,CP_-gHBfix21, and PAK, TIP4P-D^75^ with DESRES and DES-Amber, TIP3P^58^ with ROC, Chen&Garcia and BSSF1, SPC/E^76^ with OL3, and the CHARMM-modified TIP3P with CHARMM36. We used the SWM4-NDP^77^ and AMOEBA^64, 65^ water models for CHARMM_Drude_ and AMOEBA, respectively. The chosen ion parameters also differed depending on which FF and water model were utilized. For systems using the OPC, TIP3P and SPC/E water models, we used the Joung&Cheatham parameters^78^ (TIP4PEW Joung&Cheatham ion parameters were used for OPC), except the OL3_R2.7_ FF where the Li&Merz parameters were used.^79^ The DESRES and DES-Amber simulations were run with the recommended Charmm22^80^ ion parameters, Chen&Garcia with Chen&Pappu^81^ ion parameters and CHARMM36 with the CHARMM ion parameters.^82^ The CHARMM_Drude_ and AMOEBA default ion parameters were used. All the RNA FF/water/ion parameters combinations utilized are listed in Table 1.

**Table 1:**
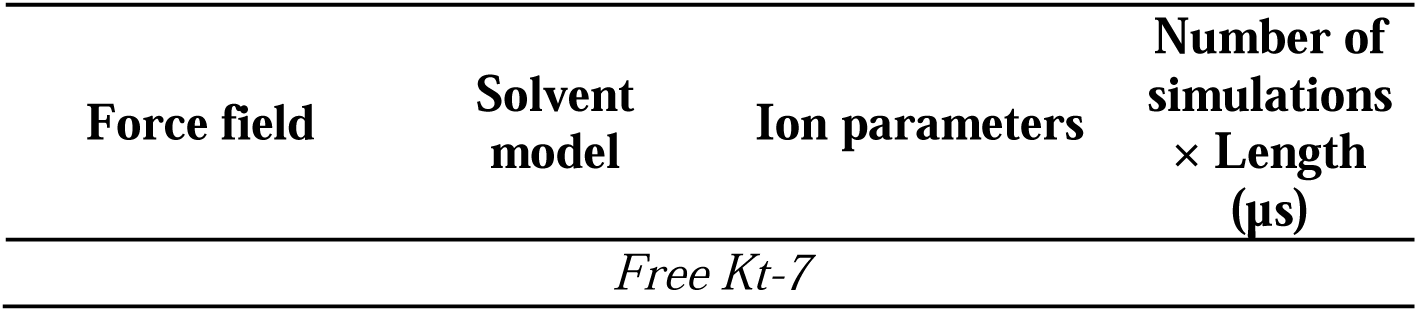
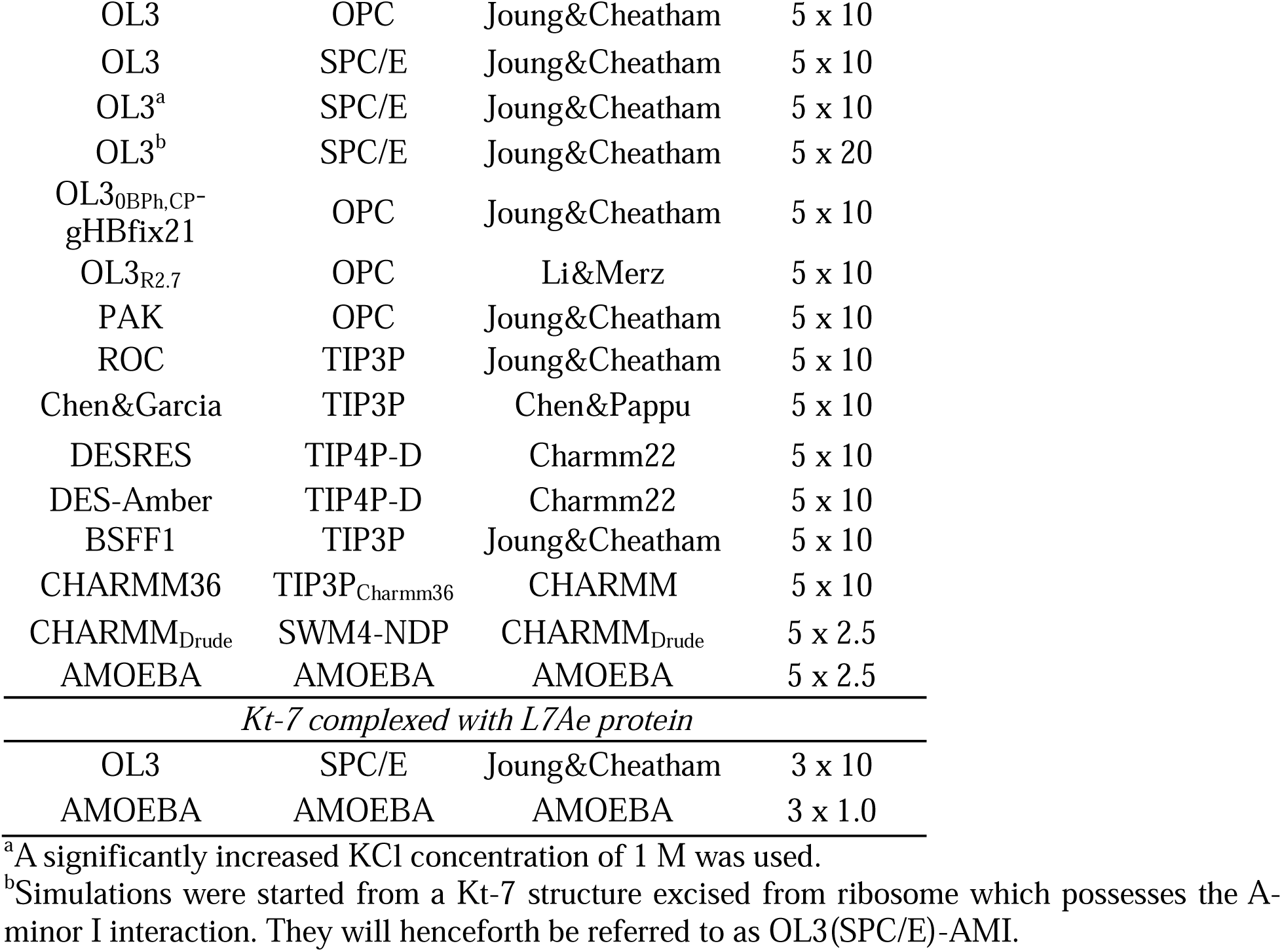
List of MD Kt-7 simulations.^a^.

### Simulation protocol – the non-polarizable FFs

The Kt-7 structure was placed in a cubic box of water molecules, with a minimal distance of 12 Å between the solute and the box border. KCl ions were added at random positions to neutralize the systems and obtain an excess-salt concentration of 0.15 M, which is well above the minimal monovalent ion concentration at which the Kt-7 was experimentally shown to autonomously fold.^24, 68^ The tLeap program of AMBER was used to generate the initial files with the exception of DES-Amber, BSSF1 and CHARMM36 FFs (see below). Excluding these FFs, the simulations were subsequently performed in AMBER20^83^ using the pmemd.MPI and pmemd.cuda^84^ programs for equilibration and production simulations, respectively. All MD simulations were run at T = 298 K, with the hydrogen mass repartitioning^85^ scheme allowing a 4-fs integration time step. Long-range electrostatics were treated with particle mesh Ewald^86^ and the distance cut-off for Lennard-Jones interactions was set to 10 Å. Production simulations were performed in a constant volume ensemble, with the temperature controlled by Langevin thermostat.^87^ For more details of the minimization and equilibration protocols, see Ref.^67^. The DES-Amber, BSSF1 and CHARMM36^59^ simulations were performed in Gromacs2020.^88^ The simulation protocol in Gromacs2020 slightly differed from the one in AMBER20 due to differences in the simulation codes. Specifically, Gromacs simulations were performed in a rhombic dodecahedral box and bonds involving hydrogens were constrained using the LINCS algorithm.^89^ The cut-off distance for the direct space summation of the electrostatic interactions was 10 Å and the simulations were performed using the stochastic velocity rescale thermostat.^90^ All production simulations with pair-additive FFs were run for 10 μs (the OL3 simulations of the Kt-7 possessing an A-minor I interaction at the start were run for 20 μs) with five independent trajectories produced (Table 1).

### Simulation protocol – the polarizable FFs

Five Kt-7 structures were pre-equilibrated using the OL3(OPC) FF and used as starting points for CHARMM_Drude_^60–63^ and AMOEBA^64, 65^ simulations. For CHARMM_Drude_, the structures were transformed into the polarizable model using the CHARMM software (version 44b1).^82^ During the conversion process, Drude particles were introduced for all heavy atoms and the lone pairs associated with each H-bond acceptor. The OPC water molecules were converted into the polarizable SWM4-NDP model.^77^ After initial minimization and equilibration procedure using the NAMD 2.13 package^91, 92^ five independent production simulations were performed at 298 K in OpenMM 8.0^93^ for 2.5 μs. Drude Langevin integrator^94, 95^ was utilized with a timestep of 1 fs. The pressure was maintained at 1 bar utilizing the Monte Carlo barostat.^96^ The covalent bonds involving hydrogens were kept rigid using the SHAKE^97^ and SETTLE^98^ algorithms for solute and waters, respectively. A constraint of 0.2 Å was applied to limit the length of Drude-nuclei bonds. Electrostatic interactions were treated using the Particle-Mesh Ewald method (PME)^86^ with a 12 Å cutoff for the real space term. Non-bonded interactions were truncated at 12 Å, using a switching function from 10 to 12 Å.

For the AMOEBA FF, pre-equilibrated Kt-7 structures were transferred into xyz coordinates by Tinker and minimized in 10 000 steps using the steepest descent method. The systems were then heated up to 298 K and equilibrated for the pressure of 1 bar. Stochastic velocity rescale thermostat^90^ and Monte Carlo barostat^96, 99^ with coupling constants of 0.1 ps were used to maintain temperature and pressure, respectively. The applied real-space cutoff for electrostatic and van der Waals interactions was 7 and 12 Å, respectively. RESPA integrator^100^ was used with the integration step of 1 and 2 fs for equilibration and production simulations, respectively. Other control functions and parameters were set to their default values. We ran five independent simulations for 2.5 μs in NVT ensemble using the GPU-accelerated 2023 Tinker-HP 1.3 code.^101^

### Simulation protocol – the L7Ae protein/Kt-7 complex simulations

The FFs with the best performance in simulations of the isolated Kt-7 (OL3(SPC/E) and AMOEBA; see Results and Discussion) were further tested in the context of its complex with the L7Ae protein. The X-ray structure of this protein-RNA complex (PDB: 4BW0) was used as the starting point.^19^ The OL3 FF for RNA was paired with the ff14SB protein FF.^102^ Pre-equilibrated structures for the AMOEBA simulations were prepared using the aforementioned OL3(SPC/E)/ff14SB FF combination. All other parts of the simulation protocol mirrored those used for the isolated Kt-7 (see above). Three trajectories were generated for each FF, with lengths of 10 μs for OL3(SPC/E)/ff14SB and 1 μs for AMOEBA. The shorter timescale of AMOEBA simulations was due to the significantly larger computational costs associated with the use of polarizable FF for a system of this size.

### Analysis of structurally essential H-bonds in Kt-7

All analyses were performed using cpptraj module of Amber 20. To evaluate stability of the kink-turn motif, we monitored the presence of key H-bonds, considering them formed when the distance between heavy atoms was below 4.0 Å and the donor-hydrogen-acceptor angle above 120°. The angle condition was not imposed on H-bonds involving the O2′-groups as it would have led to excessive false negatives due to their greater conformational flexibility. The first H-bond to be analyzed was the signature interaction (SI), G_L1_(O2’)-A_1n_(N1) (Figure 1B and 2A). In many simulations, the SI tends to reversibly fluctuate between the *native* G_L1_(O2’)-A_1n_(N1) arrangement and the G_L1_(O2’)-A_1n_(N6) H-bond, which we call *non-native* SI throughout the paper (Figure 2B). The populations of both arrangements were evaluated. In the present study we generally consider the SI as formed for both arrangements, albeit to our knowledge the latter H-bond has never been observed in kink-turn experimental structures. We nevertheless consider its reversible formation as a minor structural perturbation which does not indicate a major FF imbalance.

**Figure 2:**
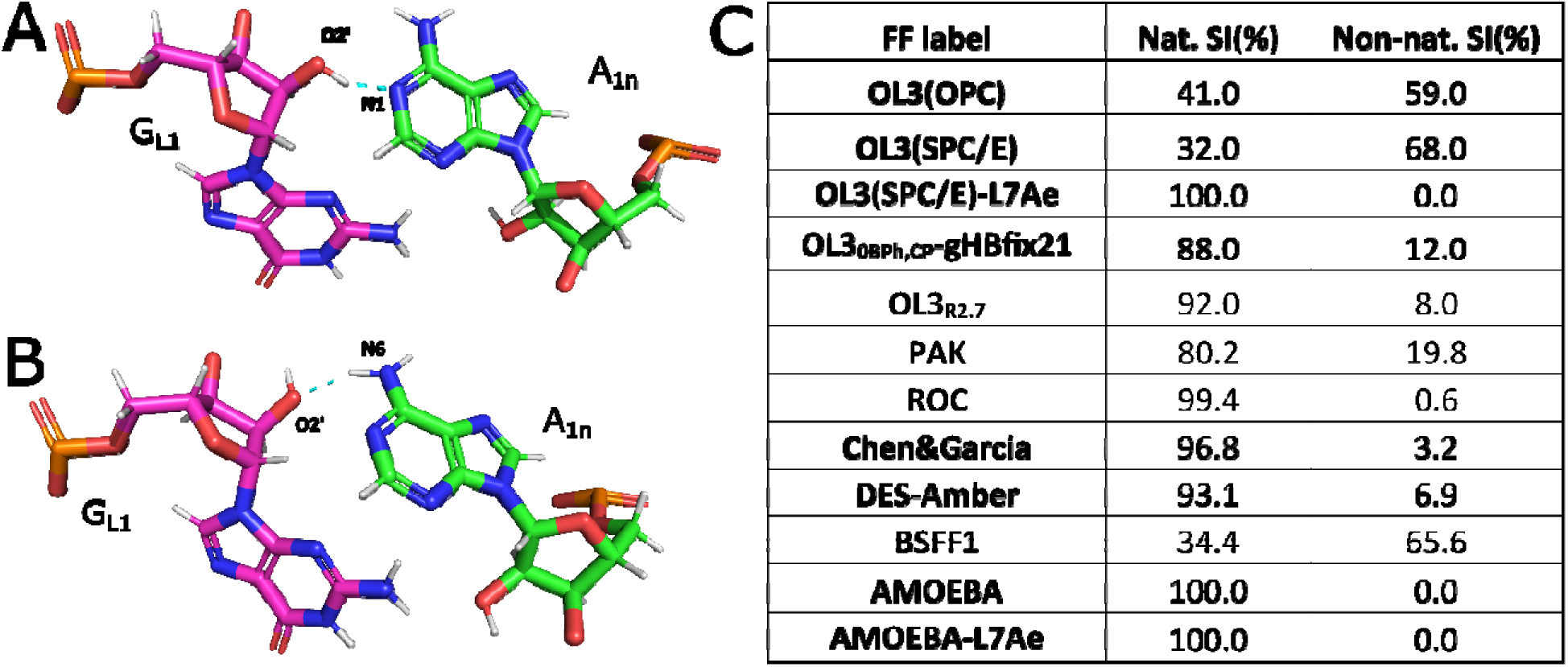
The native and non-native SI conformations as observed in MD simulations of Kt-7. A) The native and B) non-native SI. C) Table showing the normalized populations of the native and non-native SI conformations in the combined simulation ensembles. In simulations where both populations existed, they regularly exchanged on the nanosecond timescale. L7Ae indicates simulations with the bound protein discussed later.

The A-minor 0 interaction of the starting structure (PDB: 4C40) can sometimes transition into the A-minor I arrangement in the simulations. Therefore, we monitored the two sets of H-bonds corresponding to both A-minor types (Figures 1C). Specifically, G_-1n_(O2’)-A_2b_(N3) and G_-1n_(O2’)-A_2b_(O2’) H-bonds for A minor 0 referred to as AM0_A_ and AM0_B_, respectively, and G_-1n_(O2’)-A_2b_(N1), C_-1b_(O2’)-A_2b_(O2’) and A_2b_(N3)-G_-1n_(N2) H-bonds denominated as AMI_A_, AMI_B_ and AMI_C_, respectively, for A-minor I. When the A-minor I interaction is formed the simulations commonly populate temporary states with a water insertion into the C_-1b_(O2’)-A_2b_(O2’) interaction. Formation of this water bridge does not constitute disruption of the structure but rather its natural dynamics; such arrangements are in fact seen also in some experimental kink-turn structures.^103, 104^ Thus, our analyses considered the A-minor I formed even when this H-bond showed values above the cut-off, as long as the other two H-bonds were established.

The H-bonds constituting the three *t*HS AG base pairs of the NC stem, namely, the A_1n_(N6)-G_1b_(N3), G_1b_(N2)-A_1n_(N7), A_2b_(N6)-G_2n_(N3), G_2n_(N2)-A_2b_(N7), A_3b_(N6)-G_3n_(N3) and G_3n_(N2)-A_3b_(N7) (Figure 1) were also monitored. In the C-stem, we monitored H-bonds constituting the Watson-Crick base pair closest to the bulge region – G_-1n_(O6)-C_-1b_(N4), G_-1n_(N1)-C_-1b_(N3) and C_-1b_(O2)-G_-1n_(N2). This GC base pair is also involved in the A-minor interaction (Figure 1B).

### Additional analyses conditioned by the presence of SI

The following analyses were conducted exclusively on portions of the simulation ensembles where the SI remained stable, as defined above. This approach was chosen to minimize noise in the data as empirical observations showed that unstable SI essentially always implied severely disrupted kink-turn structures; the SI is the most important H-bond in the kink-turn (see the Introduction). In this fashion, we monitored the χ-glycosidic dihedral of A_L2_, with values in the range of -90° to 90° considered to be *syn* and outside of this range as *anti* conformations. Stacking of A_L2_ with either the G_L1_ or A_1n_ was evaluated by calculating the distance between geometric centers of the endocyclic N atoms of each base. Stacking was considered present for distances below 4.0 Å and 4.5 Å for the A_L2_/A_1n_ and A_L2_/G_L1_ stacks, respectively. These cut-off values were chosen empirically to well reproduce the results obtained by visual analysis. The 4BPh interaction was considered present if either the A_2b_(OP2)-G_3n_(N1) or A_2b_(OP2)-G_3n_(N2) or both H-bonds were formed (see above for general H-bond presence definition). We also evaluated the presence of the A_L3_(O2’)-A_L2_(OP1) and G_1b_(O2’)-G_2n_(N2) H-bonds, constituting the characteristic sugar-phosphate and sugar-base interactions, respectively (Figure 1B). Individual RNA acceptor atoms were considered to be binding a K^+^ cation if a K^+^ ion was located within 4 Å. The density grid showing the strongest K^+^ binding sites was visualized with VMD. Backbone dihedrals andsugar puckers,^105^ were calculated with cpptraj and visualized using python’s matplotlib module. We also monitored the correlated time development and distribution of the backbone dihedral suites^22^ exhibiting non-canonical α/γ values in the experimental kink-turn structures, i.e., the G_3n_/G_2n_, A_1n_/G_-1n_, G_L1_/A_L2_ and A_L2_/A_L3_ suites.

## Results and discussion

### General comments and the global stability of Kt-7

In the following, we describe performance of the individual FFs both in a global sense (i.e. whether the kinked shape of Kt-7 is maintained) and in the context of the individual characteristic interactions, going from the most essential to less important interactions. We have performed 91 standard MD simulations with cumulative length of 758 μs. Multiple independent trajectories were generated using each FF (Table 1), revealing a consistent performance but relatively large variability in the timescales of specific events. In other words, while characteristic transitions or instabilities consistently occurred for a given FF, the timing of their initial occurrence varied significantly. Despite this variability, we propose that the simulations were sufficiently long to evaluate the performance of each FF. Unless otherwise specified, the observations described below were identified in all or most independent trajectories obtained for a given FF. Note that we observed irreversible unkinking of Kt-7 and/or serious H-bond disruptions of the non-canonical stem in at least one trajectory using the BSSF1, DES-Amber, ROC, OL3_0BPh,CP_-gHBfix21 and OL3_R2.7_ FFs (see Supporting Information for the description of the unkinking, Table S2 and S3).

For three FFs (CHARMM36, CHARMM_Drude_ and DESRES) the simulations did not reveal a sufficiently accurate description of the kink-turn. Thus, we were not able to obtain a reliable statistics of the individually analyzed structural features due to limited sampling of the native or at least near-native kink-turn structures. To streamline the presentation, these FFs are not included in the following paragraphs analyzing the individual interactions. However, their overall performance is discussed at the end.

### Kt-7 signature interaction (SI) populates two conformations in MD simulations

The SI is the most important and defining interaction of the kink-turn motif (Figure 1B and 2).^3^ Whenever a global loss of the kink-turn’s kinked shape (see above) occurred the SI was typically the last tertiary interaction to disappear. Thus, the presence of the SI serves as a reliable and straightforward indicator of the overall structural integrity of the kink-turn. Although instances of temporary, localized disruptions of the SI without complete unfolding of the kink-turn were observed, such events were relatively rare. As observed earlier,^66, 67, 106^ many of our simulations revealed the SI sampling two distinct conformations that interchanged rapidly on the nanosecond timescale. Specifically, the native A_1n_(N1)-G_L1_(O2’) H-bond has been regularly replaced with the G_L1_(O2’)-A_1n_(N6) H-bond, henceforth referred to as the native and non-native SI, respectively (Figure 2). These transitions were always reversible on the simulation timescale with all FFs that displayed them, and they never propagated to larger perturbations of the kink-turn. Therefore, in this paper, we consider both conformations as corresponding to a stable SI. Still, an ideal FF should obviously sample dominantly the native SI arrangement.

The individual RNA FFs exhibited strikingly different propensities to sample the two SI conformations. Specifically, the OL3 FF variants (OL3(OPC) and OL3(SPC/E)), as well as BSSF1, favored the non-native SI. The population of the native SI significantly increases, without any undesirable side-effects, for the OL3_0BPh,CP_-gHBfix21 and OL3_R2.7_ modifications. The PAK, ROC, DES-Amber, and Chen&Garcia FFs also favored the native SI. Performance of the FFs favoring the native SI can at first glance look better, however, we stress that some of these FFs had significant troubles with the description of the second main interaction stabilizing the Kt-7 – the A-minor interaction (see below). We underscore the nearly 100% population of the native SI observed with both the non-polarizable ROC and the polarizable AMOEBA FF. Notably, in case of AMOEBA, the non-native SI was not observed at all (Figure 2C). This suggests the presence of the non-native SI in simulations with non-polarizable FFs might be related to the lack of polarizability. In the case of ROC, this limitation appears to be effectively compensated for through adjustments to the backbone dihedral angles, resulting in an excellent representation of the native SI.

### A-minor 0 interaction: The Achilles’ heel of some recent FFs

Essentially all tested FFs struggled to simultaneously maintain both H-bonds defining the A-minor 0 interaction (Figure 1C, Figure 3 and Supporting Information Table S4). To compare performance of the FFs, we first considered the G_-1n_(O2’)-A_2b_(N3) H-bond interaction of the A-minor 0, which was the more stable one with all FFs, and is referred to as AM0_A_. We then used the relative stability of the second interaction, G_-1n_(O2’)/A_2b_(O2’) (AM0_B_), as a tiebreaker. Based on this evaluation, the Kt-7 A-minor 0 interaction was best reproduced by the standard OL3 variants (OL3(SPC/E) and OL3(OPC)), which nearly always maintained the AM0_A_ while also preserving the AM0_B_ to a high degree (Figure 3B). The PAK, AMOEBA, and ROC FFs showed mixed results, reproducing the AM0_A_ well but struggling with AM0_B_. Finally, the BSFF1, DES-Amber, and Chen&Garcia FFs clearly struggled with both interactions. Interestingly, the OL3 variants that included the recently proposed NBfix_0BPh_,_CP_-gHBfix21 or R2.7 FF modifications performed worse for both H-bonds compared to the unmodified OL3 variants. In conclusion, the A-minor 0 interaction in Kt-7 is best reproduced by the standard OL3 while the other FFs, often trained to improve description of other RNA systems, struggle with this tertiary interaction.

**Figure 3:**
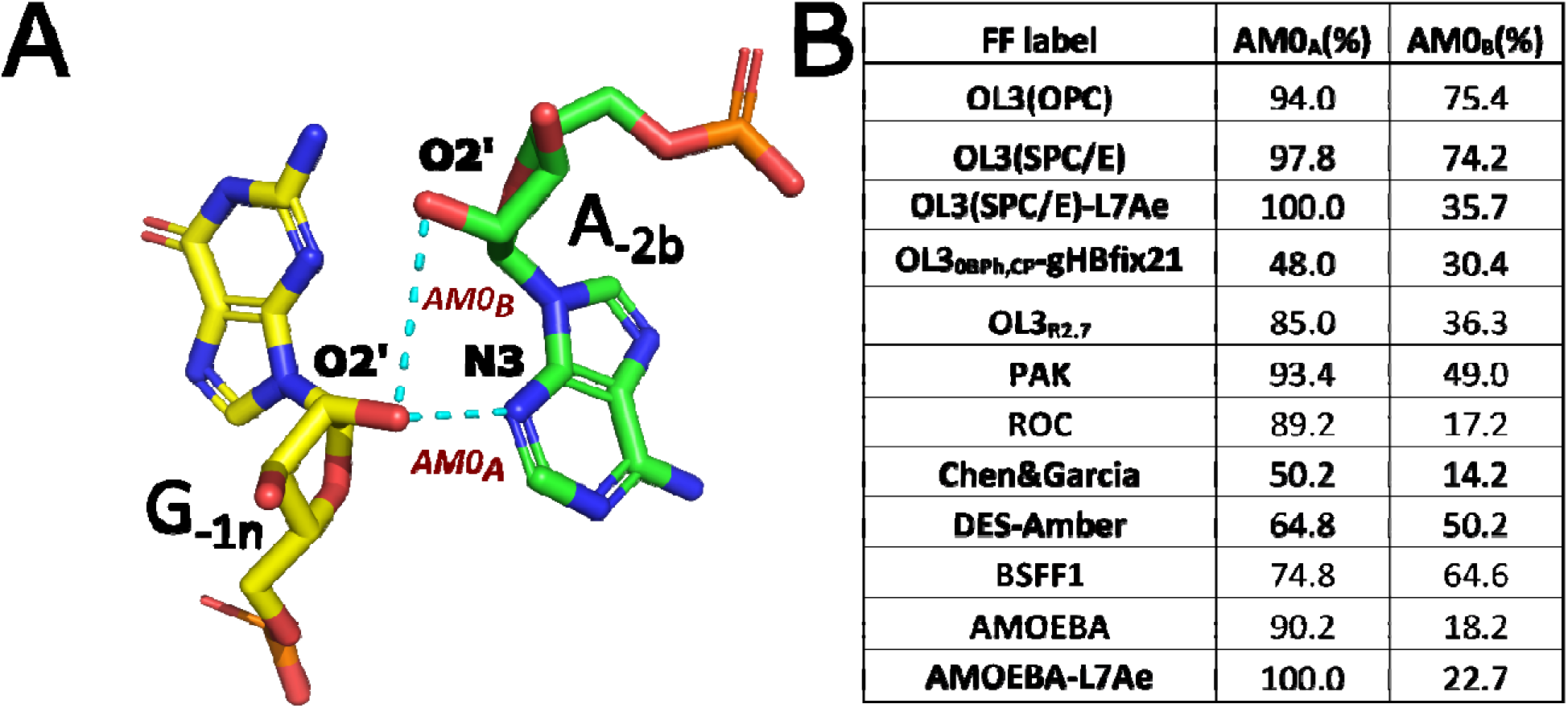
A-minor 0 interaction as observed in MD simulations of Kt-7. A) Structural overview of the interaction, with the key H-bonds indicated as AM0_A_ and AM0_B_. B) H-bond populations in full combined simulation ensembles of the individual FFs. L7Ae indicate simulations with the bound protein discussed later.

### A-minor 0 ⇆ A-minor I transitions add complexity to the simulations and their analyses

The Kt-7 structure can adopt both A-minor interaction types 0 and I (Figure 1C). While the type 0 interaction is favored in isolation and/or when the Kt-7 is bound to the L7Ae protein, the type I is observed when the Kt-7 is embedded in the ribosomal context (see also above).^19^ Consequently, if a simulation starts with Kt-7 excised from the ribosome, an eventual transition from A-minor type I to type 0 is to be expected. Indeed, previous simulation studies utilizing such starting structure universally observed A-minor transitions from type I to 0.^66, 67^ We propose that A-minor interconversion dynamics could potentially be observed also when starting from the structure of isolated Kt-7 with the A-minor 0 interaction (this study), either in sufficiently long simulations or with FFs that inherently favor the A-minor I type. While potentially interesting benchmark for the FFs, there is currently no experimental information available regarding the population balance and the expected timescale of the interconversion dynamics. Therefore, we limit our observations to noting which FFs display such A-minor transitions in our study. Namely, we regularly observed reversible A-minor transitions from 0 to I with the OL3_0BPh,CP_-gHBfix21 and Chen&Garcia FFs. In only one trajectory, a single irreversible transition from A-minor 0 to A-minor I was observed for both the AMOEBA and ROC FFs (Supporting Information Table S5). All four FFs showed problems with accurately reproducing the A-minor 0 interaction, especially the AM0_B_ (see above) H-bond. We speculate the underestimated stability of the A-minor 0 interaction could spuriously accelerate the eventual transition into A-minor I.

The A-minor transitions were not observed with other FFs in this study, which all kept the starting A-minor 0 conformation of the isolated Kt-7 (see Introduction and Methods). However, as already mentioned, when starting a simulation from the ribosomal Kt-7 structure, an eventual A-minor I to 0 transition is expected. It could serve as an additional test for the FF performance, i.e. to see whether and how the FF handles the transition. To demonstrate this, we conducted 20-μs-long simulations of Kt-7 excised from the ribosome using the best-performing pair-additive OL3(SPC/E) FF variant. In four simulations, the A-minor I successfully transitioned to A-minor 0 on a timescale ranging from 1.57 μs to 4.17 μs. No transition was observed in the fifth replicate. The transitions were irreversible, except for one replicate where the A-minor I briefly returned (between 1.57 and 1.78 μs) but then switched again to A-minor 0 (Supporting Information Table S5). The transition mechanism was the same in both directions – the adenine was sliding along the edge of the GC base pair without any other visible conformational changes in the core region of the kink-turn.

It is notable that we observed different stabilities of the other kink-turn interactions depending on the state of the A-minor interaction. For instance, for the OL3(SPC/E) FF, the native SI conformation was strongly favored in presence of A-minor I whereas the non-native SI dominated the ensemble in presence of A-minor 0. Interestingly, the FFs spontaneously sampling the A-minor I (i.e. OL3_0BPh,CP_-gHBfix21 and Chen&Garcia) didn’t show this correlation and the native SI was maintained with both A-minor types (Figure 2C and Supporting Information Table S6). The potential for A-minor transitions in Kt-7 adds an additional layer of complexity to the analysis of the FF performance, as optimally performing FF should prefer the A-minor 0 interaction for isolated Kt-7. However, the presence of some A-minor I population should not necessarily be considered a major FF issue as the genuine free energy differences between the two conformations could be relatively small.

### The base pairs of the NC- and C-stems

The GC base pairs of the C-stem and the AG base pairs in the NC-stem were generally stable in simulations with the reasonably performing FFs. In case of the AG base pairs, we observed temporary and reversible disruptions in response to fluctuations of the SI interaction. This was especially the case with the AG base pair closest to the bulge (which also acts as the acceptor for the SI) where mainly the A_1n_(N6)-G_1b_(N3) H-bond could fluctuate. Fluctuations of the N6-N3 H-bonds were observed also for the other two AG base pairs but with much lower frequency and range. The N2-N7 H-bonds were generally more structurally stable than the N6-N3 H-bonds and showed relatively larger fluctuations only with the ROC and DES-Amber FFs. In agreement with experiments and previous MD simulations,^19,107^ we have observed that formation of the A-minor I interaction (see above) is straining the geometry of the first AG base pair, making the N6-N3 H-bonds longer or disrupted.

### The A_L2_ adenine shows *syn*-*anti* transitions and alternative stacking arrangements

The second base of the bulge (A_L2_) possesses a *syn* conformation of its N-glycosidic dihedral angle in the starting structure. This seems to be the case in most kink-turns when the second bulge nucleotide is a purine, and it can be considered a characteristic feature of the kink-turn motif.^108^ When bound to the L7Ae protein, *syn*-specific protein-RNA interactions involving A_L2_ form. However, we also note that A_L2_ is the only segment of the RNA showing significant crystal packing contacts in the X-ray structure of the isolated Kt-7 (Supporting Information Figure S1) which is our starting structure. These could be artificially stabilizing the A_L2_ *syn* conformation. Indeed, in virtually all simulations using the non-polarizable FFs, the A_L2_ base consistently transitioned into the *anti* conformation (Figure 4 and Supporting Information Table S7). Only a few very short-lived returns to the *syn* state were observed and the transition can be considered essentially irreversible. The only exception was the PAK FF and polarizable AMOEBA FF which regularly allowed reversible *syn*-*anti* dynamics, however, the *anti* conformation was still preferred.

**Figure 4:**
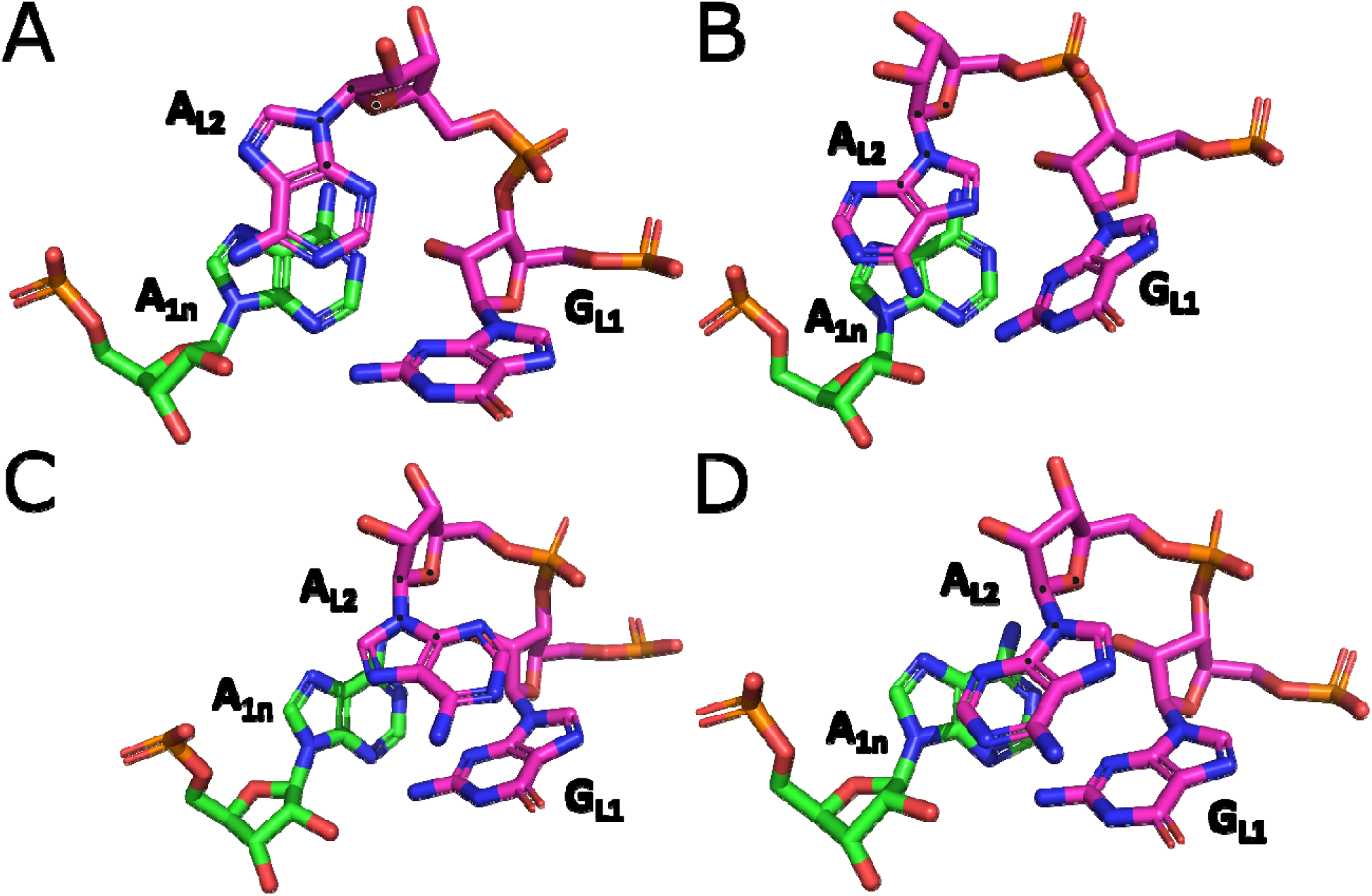
The stacking patterns and conformational states observed for the A_L2_ base. (A) A_L2_ in *syn*, stacking with A_1n_ (the native, i.e. crystal, arrangement); (B) A_L2_ in *anti*, stacking with A_1n_; (C) A_L2_ in *syn*, stacking with G_L1_; (D) A_L2_ in *anti*, stacking with G_L1_. The atoms defining the N-glycosidic dihedral angle of A_L2_ are marked with black dots.

In addition to the *syn*/*anti* transitions, the A_L2_ also showed alternative stacking patterns. Specifically, A_L2_ stacked with the A_1n_ in the starting structure while all FFs except AMOEBA preferred an alternative stacking arrangement of A_L2_ with G_L1_ (Figure 4 and Supporting Information Table S7). Formation of the alternative stacking pattern appeared unrelated to the *syn*/*anti* transitions. It cannot be ruled out that the stacking pattern may also be affected by crystal packing.

### The 4BPh and sugar-phosphate interactions disappear with most FFs

The A_2b_(OP2)-G_3n_(N1/N2) 4BPh interaction (Figure 1B) disappeared shortly after the start of simulations with all non-polarizable FFs (Supporting Information Table S7). Once lost, this interaction was rarely restored for more than a few nanoseconds. It was, however, often re-established upon formation of the A-minor I state. This suggests a potential conflict among individual interactions within the kink-turn structure. Indeed, the presence of A-minor I highly correlated with the stability of the 4BPh interaction in OL3(SPC/E)-AMI, OL3_0BPh,CP_-gHBfix21, ROC and AMOEBA simulations (Supporting Information Table S6). The AMOEBA FF maintained the 4BPh interaction in over 40% of the whole simulation ensemble despite being dominantly sampling the A-minor 0 arrangement (see above). In the experimental structures the 4BPh interaction is absent upon binding of the L7Ae protein, but it is present in the isolated Kt-7.^19^ Therefore, we consider its swift loss in the simulations a potential FF-related problem, essentially for all FFs except of AMOEBA. We hypothesized that the stability of the 4BPh interaction might be influenced by the salt conditions. Thus, we conducted OL3(SPC/E) simulations with an excess-salt concentration of 1 M (see Table 1), however, these simulations again revealed the swift loss of the 4BPh interaction. In fact, the increased salt concentration did not have any visible effect on any monitored interactions, suggesting that the salt concentration does not play a considerable role, at least not on our simulation timescale (Supporting Information Table S4, S7 and S8).

We also monitored the sugar-phosphate and sugar-base interactions native to Kt-7 – the A_L3_(O2’)-A_L2_(OP1) and G_1b_(O2’)-G_2n_(N2) H-bonds, respectively. The sugar-phosphate interaction was typically lost shortly after the simulation start for all FFs, with the notable exception of the polarizable AMOEBA FF. This loss appeared to be coupled with the A_L2_ base transitioning from the *syn* to *anti* conformation, a feature common to all tested FFs (see above). The second interaction was stable across all simulations, except for the DES-Amber and OL3_0BPh,CP_-gHBfix21 FFs. Temporary disruptions of this interaction were observed in response to changes elsewhere within the kink-turn, such as A-minor transitions or shifts in the stacking pattern of A_L2_. In general, the presence of the A-minor I and/or A_L2_/A_1n_ stacking destabilized the G_1b_(O2’)-G_2n_(N2) interaction. Once again, the exception was the polarizable AMOEBA FF which usually maintained the G_1b_(O2’)-G_2n_(N2) interaction regardless (Supporting Information Table S7).

### Binding of the L7Ae stabilizes the kink-turn’s signature interaction

To further explore the structural dynamics of Kt-7, we took the two “best-performing” FFs (OL3(SPC/E) and AMOEBA) and carried out simulations with the bound L7Ae protein (see Methods). In case of OL3(SPC/E) the binding of the protein fully stabilized the native arrangement of the SI, contrasting the simulations of the isolated Kt-7 (Figure 2C). The bound protein also affected stability of the individual A-minor 0 H-bonds (Figure 3A). The AM0_A_ H-bond was essentially fully stable with both FFs while the population of AM0_B_ interaction decreased and slightly increased in the OL3(SPC/E) and AMOEBA simulations, respectively (Figure 3B, Supporting Information Table S9).

The A_L2_ base remained in the *syn* conformation in all simulations with bound protein as it was engaged in hydrophobic contacts with the protein, blocking any potential transitions to the *anti* conformation. Surprisingly, with OL3 we observed the 4BPh interaction being more populated in simulations of the protein-RNA complex than in the isolated Kt-7, despite this interaction being present and absent in the starting structures of the isolated and bound kink-turn, respectively. With AMOEBA, we observed the opposite trend (Supporting Information Table S9). Lastly, the protein-RNA interface interactions were generally more fluctuating with the AMOEBA FF, and we observed increased dynamics of the entire protein-RNA interface. However, these fluctuations were entirely reversible, and the interface of the complex can thus be considered as fully stable in both tested FFs.

### The FFs struggle to reproduce the non-canonical _α_/_γ_ backbone dihedral suites of Kt-7

We have focused our analysis of the RNA backbone dihedrals on four sugar-phosphate backbone segments (so-called suites)^22^ exhibiting characteristic non-canonical α/γ states. These include gauche+/trans (g+/t) state for suites G_3n_/G_2n_ and A_1n_/G_-1n_, g+/g-for G_L1_/A_L2_, and g+/g+ for A_L2_/A_L3_ (Figure 5, Supporting Information Table S1). These backbone states are highly conserved for the kink-turn motif in general^22^ and their reproducibility by the FFs is another benchmarking opportunity, because the dihedral potentials are a common target in FF reparameterization efforts. For instance, the g+/t combination for α/γ has been explicitly penalized by most AMBER-derived FFs since the time of the parmbsc0 modification.^52^ Indeed, for the G_3n_/G_2n_ suite we observed an immediate and permanent transition of all tested FFs into canonical A-RNA values (g-/g+). The native values were sometimes briefly populated with the Chen&Garcia and AMOEBA FFs, but only as very minor states (Supporting Information Figures S2 and S3). In contrast, the native g+/t state of suite A_1n_/G_-1n_ was not completely eliminated, however, significant population of the non-native canonical A-RNA values were still present with most tested FFs. The best performance was seen with the OL3(SPC/E), Chen&Garcia, PAK, DES-Amber, ROC and AMOEBA FFs where only minor populations of the non-native canonical A-RNA values were observed. There is a surprising difference between the OL3(SPC/E) and OL3(OPC) simulations, indicating the potential importance of the water model choice (Supporting Information Figure S3B). For the G_L1_/A_L2_ suite, the native g+/g-combination immediately and permanently transitioned into canonical A-RNA values (g-/g+) for all tested FFs except of the AMOEBA FF and the protein-RNA complex simulations. Lastly, for the A_L2_/A_L3_ suite, the native g+/g+ combination was correctly reproduced by all tested FFs with only minor canonical A-RNA populations observed (Supporting Information Figures S2 and S3). The present results confirm earlier observations that it is challenging to describe the non-canonical α/γ states in RNA by MD.^109^

**Figure 5:**
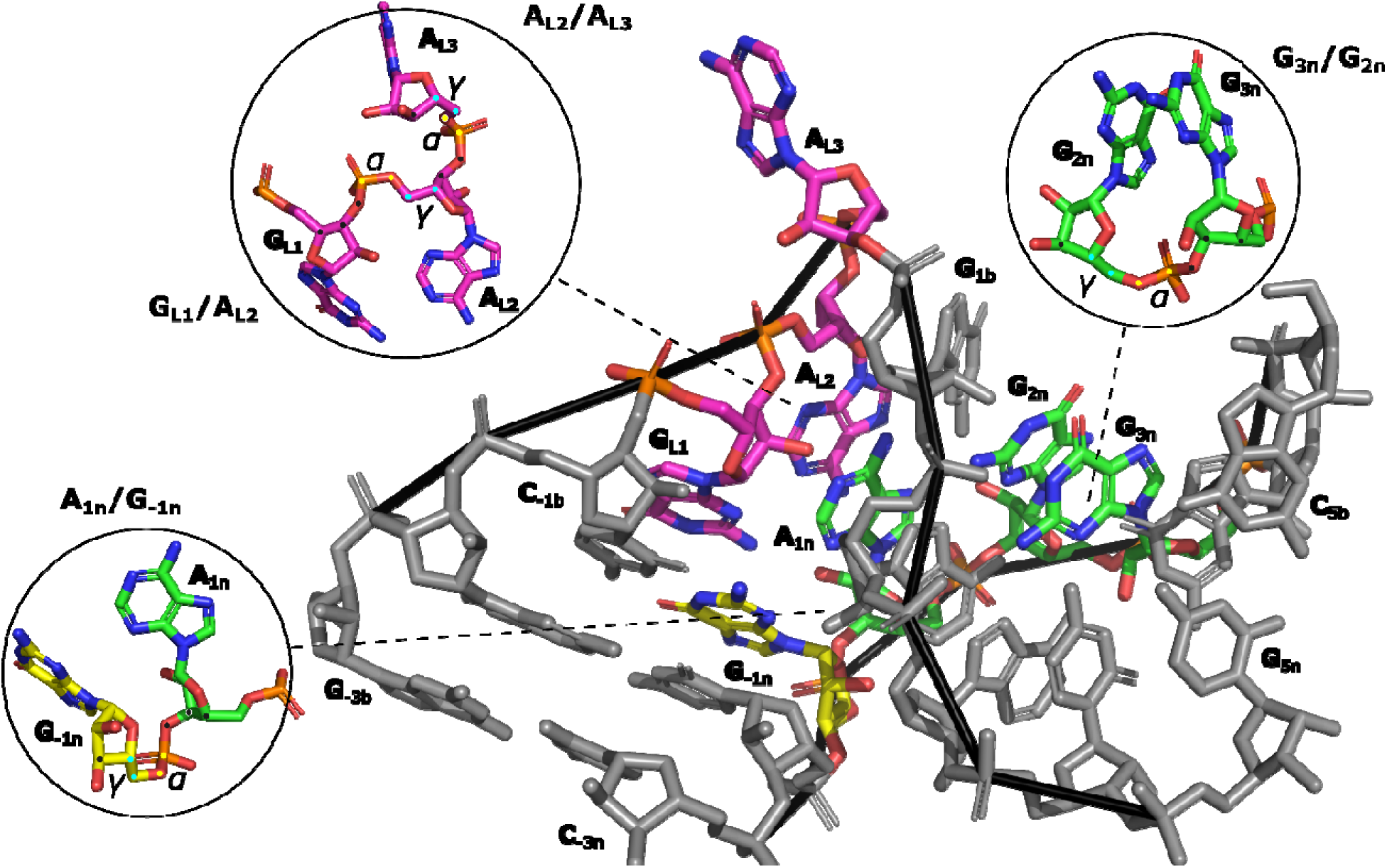
Non-canonical α/γ backbone dihedral suites of Kt-7. Kink-turn residues involved in the suites are colored as in Figure 1 while the remaining residues are grey. The insets show each suite in detail. The central atoms defining the α/γ dihedrals are colored yellow and cyan, respectively. The other atoms of the suites are indicated with a black dot.

### Additional comments on the FF performance

As noted at the beginning, we were unable to obtain a single Kt-7 trajectory using DESRES, CHARMM36, and CHARMM_DRUDE_ FFs in which the major characteristic interactions (SI and A minor 0/I) would be consistently present. As a result, these trajectories would not provide relevant statistics to assess the individual kink-turn features discussed above. Nevertheless, even these trajectories were carefully monitored and analyzed.

In case of DESRES, we observed gradual loss of the kink-turn’s characteristic interactions and often complete loss of the kinked shape (Supporting Information Table S2 and Figure S4). It is in agreement with previously conducted simulations of the Kt-7 starting from the A-minor I conformation.^67^ The simulation performance was significantly improved with the DES-Amber FF (a reparameterization of the original DESRES FF),^47^ however, DES-Amber still struggled with describing the A-minor interaction, reproducing it quite poorly compared to OL3 and most other FFs (see above and Figure 3B). This is consistent with another recent study where Kt-7 possessing an A-minor I interaction was used as starting structure.^72^ In the present work, we even observed irreversible loss of the kinked shape in one of the DES-Amber simulations (Supporting Information Table S2). It should be noted, however, that single unkinking or stem disruption events occurred also with some other FFs. Unfortunately, we know neither the correct balance between the kinked and unkinked populations, nor the kinetics of such transitions. Thus, a single unkinking event cannot be a priory interpreted as a FF failure, though it is a warning sign. Disruption of either stem, however, can be considered an FF failure, as NMR observations (for another kink-turn) point to unkinking, but not to rearrangements of base pairing or major stem disruptions.^110^

Such large structural changes occurred with the CHARMM36 FF, where we observed swift loss of the interactions characterizing the kink-turn motif after the simulation start, and shortly after that also partial or even complete disruption of both stems (Supporting Information Tables S3, S4, S8 and Figure S5). This corresponded to a loss of the entire structure, preventing us from deriving any useful information out of the CHARMM36 simulations. The problems of CHARMM36 FF to describe the stems are consistent with our recent study on the UUCG tetraloop.^72^ In contrast, the polarizable CHARMM_DRUDE_ FF showed markedly better performance compared to both CHARMM36 and DESRES, with some interactions of the Kt-7 being excellently reproduced (e.g. the 4BPh and sugar-phosphate interaction). The reason why we ultimately excluded the CHARMM_DRUDE_ simulations from our main analyses was its inability to maintain the A-minor interaction, which was universally lost soon after the trajectory start and never reformed in any trajectories except of a few short fluctuations (Supporting Information Table S4 and S8). In fact, had the A-minor interaction been maintained, the CHARMM_DRUDE_ FF would have been ranked among the better performing FFs for Kt-7. The benefit of explicit polarization is quite apparent as CHARMM_DRUDE_ successfully reproduced interactions and conformations where AMOEBA FF as well showed improvements over the non-polarizable FFs (e.g. native SI, A_L2_ in *syn* conformation, the 4BPh interaction and the sugar-phosphate interaction). Indeed, the polarizability could be important for the description of systems like Kt-7, making it a useful benchmark for refining polarizable FFs. Unfortunately, without adequately reproducing the critically important A-minor interaction, CHARMM_DRUDE_’s performance for Kt-7 cannot be deemed satisfactory at the moment. For some other recent benchmarks of the RNA CHARMM_DRUDE_ FF see refs.^72, 92, 111, 112^.

Supporting Information Table S10 summarizes the occupancies of ion binding sites across all Kt-7 simulations, revealing some notable differences among the tested FFs. Interestingly, OL3(SPC/E) simulations exhibit significantly more extensive ion binding than OL3(OPC) simulations. This highlights how the quantitative picture of ion binding is influenced by the balance of solute, water, and ion FF parameters. However, a more detailed analysis is beyond the scope of this study, as we do not have any relevant experimental information to compare with (see Supporting Information for more details).

In conclusion, among the tested FFs, the OL3(SPC/E), OL3(OPC) and AMOEBA showed the best overall performance for Kt-7. The polarizable AMOEBA was especially successful, maintaining some of the native interactions which were lost with all the other FFs. We, however, note that simulations with AMOEBA and CHARMM_DRUDE_ FFs were four times shorter compared to simulations using the non-polarizable FFs. Nevertheless, all FF problems described above tended to occur relatively early in simulations. Therefore, we suggest the good performance of AMOEBA FF is quite conclusive. From the non-polarizable FFs, we wish to emphasize the good performance of the OL3(SPC/E) and OL3(OPC) variants, i.e. the standard OL3 RNA FF, without any solute FF modifications applied (see Methods). In particular, the OL3 FF described the A-minor interaction – a frequent point of failure – best among the tested FFs. ROC and PAK were other non-polarizable FFs with an overall good performance, capable of maintaining both the native SI and the non-canonical backbone dihedrals in a satisfactory way. However, they struggled with the A-minor interaction. Table 2 presents a final qualitative overview of the performance of all tested FFs.

**Table 2:**
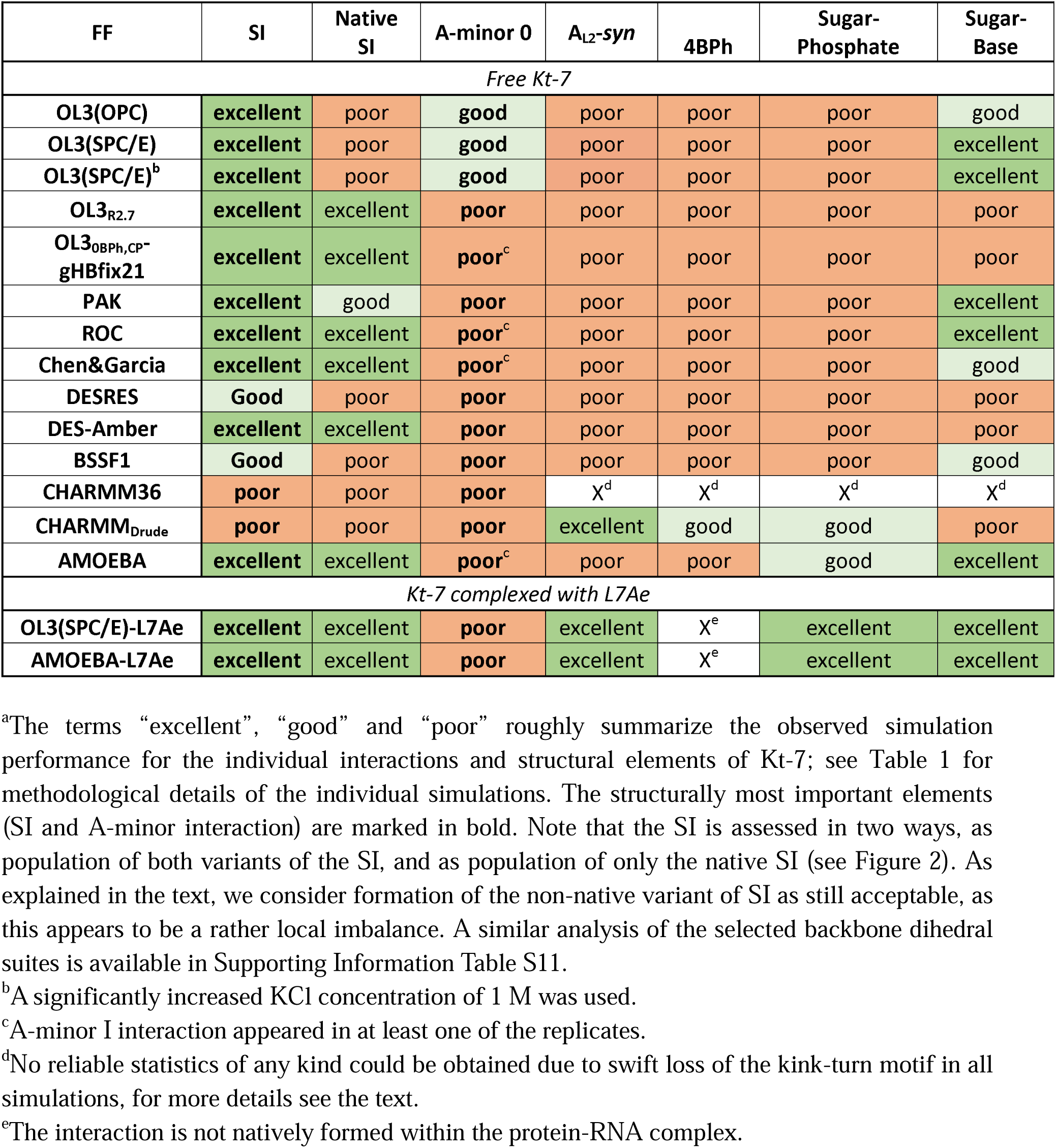
Overview of the FF performance in simulations of Kt-7.^a^.

## Conclusions

We present a large-scale benchmark MD simulation study of Kink-turn 7 (Kt-7), a prominent three-dimensional RNA motif, using a broad range of FFs. None of the tested RNA FFs provides a perfect description of Kt-7, highlighting the challenge of accurately balancing its diverse and interdependent structural features in contemporary MD simulations. Particularly concerning is the difficulty in modeling the A-minor tertiary interaction, even with recently developed FFs such as DES-Amber and OL3_0BPh,CP_-gHBfix21, which otherwise improve simulations of simpler motifs like tetranucleotides and tetraloops. This underscores the risks of overfitting FF parameters to homogeneous training sets, as more "general" FFs, such as basic OL3, ROC, or AMOEBA, often performed better. Among the tested FFs, the polarizable AMOEBA model frequently emerged as a positive outlier, suggesting that explicit treatment of polarizability may benefit systems like Kt-7. However, AMOEBA also struggled to accurately represent the A-minor interaction, particularly in comparison to OL3. To further illustrate the complexity of RNA FF development and testing, we note that both ROC and AMOEBA FFs were shown to irreversibly disrupt the native fold of the UUCG RNA tetraloop, which is a notoriously challenging system also for the basic OL3 FF.^72^ In other words, the ranking of FFs based on simulations of the UUCG tetraloop and Kt-7 differ significantly. This variation in benchmark results emphasizes the necessity of testing FFs across a diverse set of RNA motifs to substantiate claims of general improvements.

We propose that Kt-7 should serve as a key benchmark in future RNA FF parametrization efforts, as it meets several desirable criteria: (1) Computational feasibility – Its relatively small size keeps computational costs manageable, even for polarizable FFs. (2) Structural complexity – Its rich network of non-canonical interactions rigorously tests FF accuracy, particularly in A-minor interactions and tertiary H-bonding. (3) Well-defined stability metrics – The stability of its folded (kinked) structure can be extensively characterized using relatively short standard simulations. For completeness, we note that an additional test for FFs could involve modeling the free-energy landscape of Kt-7’s opening-closing dynamics. However, this would require sophisticated enhanced sampling simulations. Our preliminary attempts at such free-energy simulations revealed significant challenges in defining a suitable simulation protocol, and the results will be published separately.

In summary, the folded structure of Kt-7 represents a critically important test system. With minimal computational effort, it enables a comprehensive assessment of many aspects of RNA FFs, making it an essential benchmark for future developments in the field and a valuable complement to existing model systems. Moreover, its complexity makes it a vital regression test for avoiding overfitting and ensuring that future parameter sets remain broadly transferable across RNA architectures.

## Acknowledgement

This work has been conducted in the sustainability period of the project SYMBIT No. CZ.02.1.01/0.0/0.0/15_003/0000477 as its follow-up activity. We acknowledge the use of CESNET data storage facilities [grant number LM2018140].

