## Supporting Information for "The Kink-Turn Motif: A Powerful Test for Revealing Weaknesses in RNA Force Fields"

**Ion binding sites of Kt-7.** Due to a lack of quantitative experimental data concerning the monovalent ion binding sites of the kink-turn, we limited our analyses to their qualitative description (their positions) and population statistics. The most prominent cation binding sites were observed near the Hoogsteen edges of residues G_2n_, G_3n_ and G_4n_ (Table S10 and Figure S6), in agreement with the available kink-turn experimental structures which usually indicate magnesium binding in this region.^1^ There were only negligible variations among the positions of the ion binding sites observed between the individual FFs. Instead, the differences were in the populations, albeit still minor. Specifically, we noted a slightly lower overall ion binding propensity with the polarizable FFs. In addition, some FFs (Chen&Garcia, DES-Amber and BSSF1) preferred a different order of the most occupied ion binding sites compared to the rest of the FFs, however, the effect appeared to be rather limited (Table S10).

Adding the L7Ae protein generally lowered the K^+^ binding at the G_2n_(O6) and G_3n_(O6) sites in case of the OL3(SPC/E) simulations. We suggest this to be the result of competition between the Lys56 side chain and the cation binding as Lys56 is involved in transient H-bond formation with G_2n_(N7) (total population 19.7 %). Even when the H-bond was not formed, the lysine side-chain was still present nearby, influencing the electrostatic interactions in this region and likely repelling the cations (Supporting Information Table S12 and S13). Interestingly, this effect was far less pronounced with the polarized AMOEBA FF where there were many instances of both the lysine side chain and K^+^ binding to the G_2n_(O6) atom simultaneously. Such simultaneous binding was never observed with OL3 and likely reflected the explicit inclusion of polarization of the K^+^-G_2n_(O6) and lysine-G_2n_(O6) pairs by the AMOEBA FF.

**The non-canonical C2′-endo sugar puckers.** In the main text, we discussed several backbone suites of Kt-7 possessing non-canonical α/γ backbone dihedrals. Related to these is the presence of native C2′-endo puckers in several nucleotides from or near those suites, namely A_1n_, A_L2_, A_L3_ and G_2n_. These C2′-endo puckers were well-maintained in simulations with the notable exception of A_L2_ (part of the loop region) which always transitioned into C3’-endo with all FFs except AMOEBA (Figure S7). For the sake of completeness, we also note that the native C3′-endo puckers (all other nucleotides except those noted above) were universally well reproduced by all FFs.

**Unkinking of the kink-turn and the additional disruptions which may indicate FF problems.** In the main text, the term *unkinking* refers to the loss of the SI and A‑minor interactions, followed by straightening of the Kt-7 structure, as shown in Figure S4. All the unkinking events observed in our simulations are listed in Table S2. Although unkinking is irreversible in our simulations, it does not necessarily imply a FF problem if it occurs in just one replicate (R) simulation for a given FF. The kink-turn obviously must exist in some equilibrium with its unkinked conformation and the timescale of our simulations might simply not be sufficient to capture the reverse process.

However, in vast majority of cases, we observed that the unkinking leads to additional structural changes, some of them rather severe. These are likely already reflecting FF problems. The unkinked structured of the spliceosomal U4 RNA kink-turn (Kt-U4) was solved by NMR.^2^ The Kt-U4 is, in contrast to Kt-7, unkinked in presence of monovalent ions. The unkinked conformation does not show any other structural changes besides mere unkinking of the two stems. In contrast, the only two simulations with unkinking where no other structural changes occurred was ROC (R1) and DESRES (R3). However, in case of the latter this was mostly due to the unkinking occurring at the very end of the simulation which left no time for other changes. Other DESRES replicates where unkinking occurred more swiftly revealed severe disruptions (Table S2 and S3). A minor additional change after unkinking occurred in the CHARMM_DRUDE_ (R1), which showed the disruption of the first AG base pair (A_1n_–G_1b_). This base pair was previously identified as highly sensitive to fluctuations of the SI and its disruption during unkinking could be a genuine structural effect. In addition, the CHARMM_DRUDE_ FF is unable to correctly describe the A-minor interaction (see the main text), which could also be affecting the unkinking process.

More serious changes occurred in OL3_0BPh,CP‑_gHBfix21 (R2), DESRES (R2, R5), and DES‑Amber (R4) simulations. In these simulations the A_1n_–G_1b_ was disrupted shortly after unkinking. It was then replaced by a spurious, non‑native AG base pair formed between A_1n_ and G_L1_ (Figure S8). This prevented reformation of the native AG base pair and almost certainly precluded any chance for restoring the kink-turn even on a hypothetically longer timescale as the critical SI interaction could not reform under such circumstances. This simulation development is also not supported by any experimental data.

Lastly, we observed very serious degradation of the kink-turn structure in the OL3R2.7 (R4) and DESRES (R1) simulations, where we evidenced disruption of both the Hoogsteen and Watson–Crick H-bonds of the non‑canonical stem. Such large-scale loss of the structure on relatively short timescales should most likely be considered FF issues. For the sake of completeness, we also note that with the CHARMM36 FF, both stems began losing cohesion swiftly after unkinking (Figure S5). Finally, in BSSF (R1) and DESRES (R4), disruption of the stems occurred even without any unkinking. All the stem disruption events are listed in Table S3.

**Supporting Information Tables**

Table S1: **Backbone dihedral angles and pucker observed in the Kt-7 molecule excised from its X-ray structure (PDB: 4C40).^a^**

| **Residue / Dihedral** | **Alpha(α)** | **Beta(β)** | **Gamma(γ)** | **Delta(δ)** | **Epsilon(ε)** | **Zeta(ζ)** | **Pucker^3^** |
| --- | --- | --- | --- | --- | --- | --- | --- |
| **G_5n_** | - | - | 55.6 | 76.2 | -144.0 | -83.3 | 15.1 |
| **G_4n_** | -50.0 | 163.6 | 54.5 | 79.4 | -142.0 | -59.7 | 17.2 |
| **G_3n_** | -56.9 | 172.8 | 54.2 | 84.8 | -136.0 | -120.0 | 20.3 |
| **G_2n_** | **139.1** | -120.0 | **177.4** | 148.9 | -110.2 | -175.8 | 167.5 |
| **A_1n_** | -66.8 | 163.8 | 32.6 | 149.6 | -85.2 | 101.2 | 162.9 |
| **G_-1n_** | **82.7** | -108.0 | **-179.2** | 84.1 | -133.1 | -87.9 | 3.2 |
| **C_-2n_** | -37.8 | 160.4 | 42.5 | 81.6 | -147.1 | -74.6 | 18.9 |
| **C_-3n_** | -63.2 | 173.8 | 56.4 | 79.9 | - | - | 21.3 |
| **G_-3b_** | - | - | 82.8 | 85.4 | -147.7 | -72.7 | 10.7 |
| **G_-2b_** | -76.2 | -177.1 | 58.8 | 78.4 | -145.0 | -63.9 | 11.0 |
| **C_-1b_** | -68.3 | 179.9 | 59.0 | 76.7 | -145.8 | -62.4 | 23.6 |
| **G_L1_** | -64.2 | 173.9 | 50.8 | 73.8 | -126.8 | -132.4 | 21.4 |
| **A_L2_** | **46.5** | -179.5 | **-73.7** | 153.1 | -127.7 | 80.5 | 165.4 |
| **A_L3_** | **74.3** | -174.6 | 59.7 | 150.0 | -86.8 | -41.4 | 174.2 |
| **G_1b_** | -82.2 | -151.0 | 68.6 | 153.3 | -136.8 | 146.1 | 167.0 |
| **A_2b_** | -71.8 | 174.0 | 42.0 | 90.4 | -136.9 | -73.1 | 0.4 |
| **A_3b_** | -70.1 | 164.9 | 65.4 | 80.6 | -132.4 | -62.5 | 19.4 |
| **C_4b_** | -65.7 | 170.1 | 52.4 | 80.8 | -148.9 | -74.2 | 12.1 |
| **C_5b_** | -56.4 | -178.0 | 43.6 | 78.7 | - | - | 13.3 |

^a^The α and γ values deviating from the canonical A-RNA backbone conformation are in bold.

Table S2: **Irreversible unkinking of Kt-7 observed in MD simulations.**

| **FF label** | **Unkinking (µs)**^a^ |
| --- | --- |
| **OL3_0BPh,CP_-gHBfix21** | R2(3.85)^bc^ |
| **OL3_R2.7_** | R4(3.44)^b^ |
| **ROC** | R2(8.48) |
| **DESRES** | R1(0.70)^bc^,R3(9.24),R5(6.71)^bc^ |
| **DES-Amber** | R4(6.96)^bc^ |
| **CHARMM36** | R1,R2,R3,R4,R5^b^(all shortly after start) |
| **CHARMM_Drude_** | R1(0.77)^b^ |

^a^Indicates the simulation time (µs) and replicate in which the unkinking occurred. The RNA FFs not stated (see main text Table 1) showed no instances of unkinking.

^b^Disruption of the first AG base pair (A_1n_-G_1b_).

^c^Disruption of the first AG base pair (A_1n_-G_1b_), followed by formation of non-native and spurious AG base pair between A_1n_ and G_L1_.

Table S3: **Irreversible disruptions of Kt-7’s stems observed in MD simulations.**

| **FF label** | **Disruption of H-bonds (µs)^a^** |
| --- | --- |
| **OL3_R2.7_** | R4(4.11) |
| **DESRES** | R1(4.49),R4(5.23) |
| **BSSF1** | R1(2.25) |
| **CHARMM36** | R1(1.30),R2(0.17),R3(0.41),R4(0.27),R5(5.86) |

^a^Indicates the replicate and simulation time (µs) at which a subsequently irreversible disruption of a stem or stems first started to appear. Reversible fluctuations are not considered.

Table S4: **Population analyses (in %) of the characteristic H-bond interactions forming the A-minor 0 and A-minor I interactions. ^a^**

| **FF label / Interaction** | **AM0_A_** | **AM0_B_** | **AMI_A_** | **AMI_B_** | **AMI_C_** |
| --- | --- | --- | --- | --- | --- |
| **OL3(OPC)** | 94.0 (± 3.2) | 75.4 (± 10.4) | 0.2 (± 0.4) | 12.0 (± 21.6) | 0.0 (± 0.0) |
| **OL3(SPC/E)** | 97.8 (± 1.6) | 74.2 (± 4.5) | 0.0 (± 0.0) | 1.2 (± 1.5) | 0.0 (± 0.0) |
| **OL3(SPC/E)-1M^b^** | 99.6 (± 0.5) | 72.8 (± 4.7) | 0.0 (± 0.0) | 1.6 (± 1.4) | 0.0 (± 0.0) |
| **OL3(SPC/E)-L7Ae** | 100.0 (± 0.0) | 35.7 (± 2.4) | 0.0 (± 0.0) | 0.7 (± 0.5) | 0.0 (± 0.0) |
| **OL3_R2.7_** | 85.0 (± 5.9) | 36.3 (± 2.5) | 0.0 (± 0.0) | 5.3 (± 3.9) | 0.0 (± 0.0) |
| **OL3_0BPh,CP_-gHBfix21** | 48.0 (± 7.9) | 30.4 (± 7.7) | 27.6 (± 11.3) | 4.8 (± 2.7) | 22.2 (± 8.1) |
| **PAK** | 93.4 (± 7.7) | 49.0 (± 4.9) | 0.2 (± 0.4) | 1.8 (± 1.0) | 0.2 (± 0.4) |
| **ROC** | 89.2 (± 19.7) | 17.2 (± 4.8) | 12.2 (± 19.4) | 9.2 (± 13.6) | 10.0 (± 19.5) |
| **Chen&Garcia** | 50.2 (± 32.1) | 14.2 (± 11.3) | 40.4 (± 34.1) | 35.2 (± 29.6) | 39.6 (± 33.9) |
| **DESRES** | 49.6 (± 9.9) | 32.0 (± 11.4) | 20.2 (± 14.3) | 8.2 (± 9.7) | 1.8 (± 1.5) |
| **DES-Amber** | 64.8 (± 8.9) | 50.2 (± 15.0) | 4.8 (± 8.1) | 10.2 (± 10.3) | 1.8 (± 2.2) |
| **BSSF1** | 74.8 (± 30.6) | 64.6 (± 21.5) | 1.2 (± 1.6) | 15.4 (± 23.2) | 0.4 (± 0.8) |
| **CHARMM36** | 0.0 (± 0.0) | 0.0 (± 0.0) | 0.0 (± 0.0) | 0.0 (± 0.0) | 0.0 (± 0.0) |
| **CHARMM_Drude_** | 19.0 (± 11.7) | 1.2 (± 1.6) | 16.6 (± 7.9) | 9.2 (± 17.9) | 0.4 (± 0.8) |
| **AMOEBA** | 90.4 (± 17.7) | 19.2 (± 4.8) | 9.0 (± 18.0) | 1.2 (± 1.9) | 8.2 (± 16.4) |
| **AMOEBA-L7Ae** | 100.0 (± 0.0) | 22.7 (± 3.1) | 0.0 (± 0.0) | 0.0 (± 0.0) | 0.0 (± 0.0) |

^a^The values refer to combined full simulation ensembles of the individual FFs. The standard deviation values refer to the variability among the individual replicates that make up the combined ensembles.

^b^A significantly increased KCl concentration of 1 M was used.

Table S5: **Time intervals (in µs) in which the A-minor I interaction was present in the individual simulations.^a^**

| **FF label / Replicate** | **R1** | **R2** | **R3** | **R4** | **R5** |
| --- | --- | --- | --- | --- | --- |
| **OL3(SPC/E)-AMI** | 0-2.31 | 0-3.49 | 0-20 | 0-4.17 | 0-1.57;1.78-2.44 |
| **OL3_0BPh,CP_-gHBfix21** | 2.07-4.32;4.83-5.84 | 0.94-1.62;2.64-3.85 | 6.84-7.61 | 0.58-3.24;9.77-9.88 | 5.45-5.74;7.50-10 |
| **Chen&Garcia** | 8.25-10 | 1.5-10 | 4.59-6.59;9.99-10 | 2.05-6.64;6.94-10 | x |
| **ROC** | X | x | x | x | 5.05-10 |
| **AMOEBA** | X | x | x | 1.37-2.5 | x |

^a^Replicates marked with “x” did not populate the A-minor I interaction. Note that the OL3(SPC/E)-AMI simulations were started from a structure with the A-minor I interaction already present (see main text Methods). FFs not included did not sample the A-minor I interaction.

Table S6: **Population analyses (in %) of the characteristic Kt-7 interactions whenever A-minor I interaction was present.^a^**

| **Interaction / FF label** | **OL3(SPC/E)-AMI** | **Chen&Garcia** | **OL3_0BPh,CP_-gHBfix21** | **ROC** | **AMOEBA** |
| --- | --- | --- | --- | --- | --- |
| **SI** | 99.0 | 100.0 | 98.0 | 99.0 | 100.0 |
| **Native SI** | 85.0 | 92.0 | 98.0 | 95.0 | 100.0 |
| **Non-native SI** | 13.8 | 7.0 | 0.0 | 4.0 | 0.0 |
| ***syn*** | 52.2 | 0.0 | 1.0 | 4.0 | 0.0 |
| **A_L2_/A_1n_** | 46.0 | 38.0 | 4.0 | 14.0 | 0.0 |
| **A_L2_/G_L1_** | 25.4 | 41.0 | 80.0 | 60.0 | 95.0 |
| **4BPh** | 79.2 | 17.0 | 26.0 | 51.0 | 77.0 |
| **Sugar-Phosphate** | 80.6 | 20.0 | 8.0 | 42.0 | 71.0 |
| **Sugar-Base** | 81.2 | 63.0 | 70.0 | 38.0 | 53.0 |

^a^The Table shows population of the selected structural features during those parts of trajectories where the Kt-7 adopted the A-minor I conformation. Note that the OL3(SPC/E)-AMI simulations were started from a structure with the A-minor I interaction already present (see main text Methods). For the remaining FFs, the A-minor I interaction was sometimes populated in simulations starting from the A-minor 0 conformation. The Table mainly illustrates the positive correlation between the native SI (see the main text Figure 3) and the A-minor I interaction.

Table S7: **Population analyses (in %) of the selected tertiary H-bonds of Kt-7, the N-glycosidic dihedral of A_L2_ and its stacking patterns.^a^**

| **FF label / Interaction** | **4BPh** | **Sugar-Phosphate** | **Sugar-Base** | ***A_L2_- syn*** | **A_L2_/A_1n_** | **A_L2_/G_L1_** |
| --- | --- | --- | --- | --- | --- | --- |
| **OL3(OPC)** | 2.2 (± 1.3) | 33.6 (± 3.9) | 69.2 (± 35.1) | 8.0 (± 5.3) | 19.6 (± 26.7) | 56.6 (± 25.4) |
| **OL3(SPC/E)** | 5.2 (± 3.0) | 37.0 (± 3.8) | 92.6 (± 4.5) | 10.4 (± 9.3) | 8.4 (± 5.3) | 63.4 (± 6.5) |
| **OL3(SPC/E)-1M^b^** | 6.0 (± 2.8) | 42.2 (± 3.5) | 99.0 (± 0.9) | 17.0 (± 11.6) | 6.8 (± 5.7) | 65.0 (± 6.1) |
| **OL3(SPC/E)-L7Ae** | 20.0 (± 0.8) | 100.0 (± 0.0) | 100.0 (± 0.0) | 100.0 (± 0.0) | 84.3 (± 0.5) | 0.0 (± 0.0) |
| **OL3_R2.7_** | 3.3 (± 0.5) | 30.0 (± 5.4) | 58.7 (± 12.5) | 3.7 (± 0.5) | 19.7 (± 4.0) | 61.0 (± 5.7) |
| **OL3_0BPh,CP_-gHBfix21** | 7.0 (± 4.5) | 8.8 (± 4.8) | 23.6 (± 11.1) | 3.0 (± 1.3) | 19.8 (± 2.8) | 53.4 (± 4.8) |
| **PAK** | 18.4 (± 8.7) | 38.0 (± 11.1) | 84.8 (± 18.0) | 27.6 (± 16.5) | 20.8 (± 9.4) | 57.2 (± 9.2) |
| **ROC** | 8.8 (± 9.2) | 13.2 (± 7.0) | 93.8 (± 12.4) | 2.2 (± 0.4) | 2.0 (± 2.6) | 67.0 (± 2.1) |
| **Chen&Garcia** | 13.8 (± 6.4) | 13.8 (± 8.5) | 61.2 (± 20.4) | 0.4 (± 0.5) | 20.4 (± 22.7) | 47.8 (± 17.6) |
| **DES-Amber** | 1.4 (± 0.5) | 39.6 (± 10.9) | 14.0 (± 6.6) | 2.6 (± 2.1) | 28.0 (± 9.9) | 48.6 (± 13.4) |
| **BSSF1** | 3.0 (± 2.3) | 33.8 (± 3.1) | 64.2 (± 33.7) | 9.2 (± 6.2) | 17.6 (± 28.3) | 61.6 (± 30.6) |
| **AMOEBA** | 47.2 (± 9.5) | 74.0 (± 6.0) | 95.4 (± 8.2) | 42.4 (± 29.8) | 55.8 (± 24.8) | 22.4 (± 27.6) |
| **AMOEBA-L7Ae** | 27.3 (± 4.7) | 99.0 (± 0.0) | 99.7 (± 0.5) | 100.0 (± 0.0) | 89.0 (± 1.4) | 0.0 (± 0.0) |

^a^The values refer to combined full simulation ensembles of the individual FFs when the signature interaction was present. The standard deviation values refer to the variability among the individual replicates that make up the combined ensembles. Note that the A_L2_ nucleotide is involved in extensive crystal packing interactions (see the main text and Figure S1).

^b^A significantly increased KCl concentration of 1 M was used.

Table S8: **Population analyses (in %) of the signature interaction (SI).^a^**

| **FF label** | **SI** |
| --- | --- |
| **OL3(OPC)** | 90.6 (± 2.4) |
| **OL3(SPC/E)** | 92.8 (± 1.2) |
| **OL3(SPC/E)-1M^b^** | 93.8 (± 0.4) |
| **OL3(SPC/E)-L7Ae** | 100.0 (± 0.0) |
| **OL3_R2.7_** | 91.7 (± 1.7) |
| **OL3_0BPh,CP_-gHBfix21** | 92.4 (± 9.7) |
| **PAK** | 91.8 (± 1.9) |
| **ROC** | 97.4 (± 4.7) |
| **Chen&Garcia** | 98.4 (± 0.8) |
| **DESRES** | 73.6 (± 10.0) |
| **DES-Amber** | 88.8 (± 3.5) |
| **BSSF1** | 78.2 (± 28.8) |
| **CHARMM36** | 10.0 (± 12.0) |
| **CHARMM_Drude_** | 30.8 (± 7.7) |
| **AMOEBA** | 98.8 (± 1.5) |
| **AMOEBA-L7Ae** | 100.0 (± 0.0) |

^a^The values refer to combined average of simulation ensembles of the individual FFs. The standard deviation values (in parentheses) refer to the variability among the individual replicates that make up the combined ensembles.

^b^A significantly increased KCl concentration of 1 M was used.

Table S9: **Population analyses (in %) of tertiary RNA-RNA interactions in L7Ae/Kt-7 protein-RNA complex simulations and their comparison with the free Kt-7.^a^**

| **Interaction / FF Label** | **OL3(SPC/E)** | **OL3(SPCE)-L7Ae** | **AMOEBA** | **AMOEBA-L7Ae** |
| --- | --- | --- | --- | --- |
| **SI** | 92.8 (± 1.2) | 100.0 (± 0.0) | 98.6 (± 1.4) | 100.0 (± 0.0) |
| **Native SI** | 30.0 (± 6.0) | 99.3 (± 0.5) | 98.6 (± 1.4) | 100.0 (± 0.0) |
| **Non-native SI** | 63.0 (± 6.0) | 0.0 (± 0.0) | 0.0 (± 0.0) | 0.0 (± 0.0) |
| **AM0_A_** | 97.8 (± 1.6) | 100.0 (± 0.0) | 90.2 (± 17.6) | 100.0 (± 0.0) |
| **AM0_B_** | 74.2 (± 4.5) | 35.7 (± 2.4) | 18.2 (± 4.8) | 22.7 (± 3.1) |
| **AMI_A_** | 0.0 (± 0.0) | 0.0 (± 0.0) | 9.0 (± 18.0) | 0.0 (± 0.0) |
| **AMI_B_** | 1.2 (± 1.5) | 0.7 (± 0.5) | 1.4 (± 2.3) | 0.0 (± 0.0) |
| **AMI_C_** | 0.0 (± 0.0) | 0.0 (± 0.0) | 0.0 (± 0.0) | 0.0 (± 0.0) |
| ***syn*** | 10.4 (± 9.3) | 100.0 (± 0.0) | 32.6 (± 24.8) | 100.0 (± 0.0) |
| **A_L2_/A_1n_ stacking** | 8.4 (± 5.3) | 84.3 (± 0.5) | 52.6 (± 23.3) | 89.0 (± 1.4) |
| **A_L2_/G_L1_ stacking** | 63.4 (± 6.5) | 0.0 (± 0.0) | 24.0 (± 26.2) | 0.0 (± 0.0) |
| **4BPh** | 5.2 (± 3.0) | 20.0 (± 0.8) | 46.0 (± 9.7) | 27.3 (± 4.7) |
| **Sugar-Phosphate** | 37.0 (± 3.8) | 100.0 (± 0.0) | 74.6 (± 5.9) | 99.0 (± 0.0) |
| **Sugar-Base** | 92.6 (± 4.5) | 100.0 (± 0.0) | 95.2 (± 8.6) | 99.7 (± 0.5) |

^a^The values refer to combined simulation ensembles of the individual FFs. The standard deviation values refer to the variability among the individual replicates that make up the combined ensembles.

Table S10: **Potassium binding-sites population analyses (in %).^a^**

| **OL3(OPC)** | | **OL3(SPC/E)** | | **OL3_0BPh,CP_-gHBfix21** | | **OL3(SPC/E)-1M** | | **DES-Amber** | |
| --- | --- | --- | --- | --- | --- | --- | --- | --- | --- |
| **Acceptor** | **Population(%)** | **Acceptor** | **Population(%)** | **Acceptor** | **Population(%)** | **Acceptor** | **Population(%)** | **Acceptor** | **Population(%)** |
| [G_2n_(O6)](mailto:G_4@O6) | 124.4 | G_2n_(O6) | 150.3 | G_2n_(O6) | 135.6 | G_3n_(O6) | 116.4 | G_3n_(N7) | 59.5 |
| G_3n_(O6) | 103.4 | G_3n_(O6) | 145.8 | G_3n_(O6) | 132.0 | G_2n_(O6) | 115.9 | G_2n_(O6) | 58.8 |
| G_3n_(N7) | 75.7 | G_3n_(N7) | 84.3 | G_3n_(N7) | 83.1 | G_3n_(N7) | 66.9 | G_3n_(O6) | 57.0 |
|  |  | G_4n_(N7) | 79.5 | G_4n_(OP2) | 70.9 | G_4n_(N7) | 66.4 | **AMOEBA** | |
|  |  | G_4n_(OP2) | 67.4 |  |  | G_5n_(O6) | 58.2 | **Acceptor** | **Population(%)** |
|  |  | G_5n_(O6) | 58.5 |  |  | G_4n_(OP2) | 58.0 | G_3n_(O6) | 90.4 |
|  |  | G_4n_(O6) | 58.1 |  |  |  |  | G_2n_(O6) | 64.1 |
|  |  | G_1b_(N7) | 56.1 |  |  |  |  | G_4n_(OP2) | 50.0 |
|  |  | G_2n_(N7) | 51.2 |  |  |  |  | G_3n_(N7) | 50.0 |
| **OL3_R2.7_** | | **PAK** | | **ROC** | | **Chen&Garcia** | | **BSFF1** | |
| **Acceptor** | **Population(%)** | **Acceptor** | **Population(%)** | **Acceptor** | **Population(%)** | **Acceptor** | **Population(%)** | **Acceptor** | **Population(%)** |
| G_2n_(O6) | 126.3 | G_2n_(O6) | 121.9 | G_2n_(O6) | 115.8 | G_3n_(O6) | 87.1 | G_2n_(O6) | 127.3 |
| G_3n_(O6) | 107.4 | G_3n_(O6) | 104.0 | G_3n_(O6) | 100.6 | G_2n_(O6) | 84.3 | G_3n_(O6) | 122.3 |
| G_3n_(N7) | 73.7 | G_3n_(N7) | 78.9 | G_3n_(N7) | 69.2 | G_3n_(N7) | 74.8 | G_3n_(N7) | 92.7 |
| G_4n_(OP2) | 67.5 | G_4n_(OP2) | 53.1 | G_4n_(OP2) | 57.6 | G_4n_(N7) | 53.6 | G_4n_(OP2) | 77.1 |
| G_1b_(N7) | 53.4 | G_1b_(N7) | 52.1 | G_1b_(N7) | 55.6 |  |  | G_4n_(N7) | 61.5 |
|  |  |  |  | G_4n_(N7) | 52.6 |  |  |  |  |

^a^Ion-binding sites with a population above 50% are listed. Populations of over 100% indicate more than one potassium ion being present at the site on average. This analysis was done over the combined simulation ensemble of all trajectories when the SI interaction was present.

Table S11: **FF** **performance evaluation of the non-canonical** **α/****γ dihedrals from selected backbone dihedral suites of Kt-7.^a^**

| **FF** | **α-G_3n_/G_2n_** | **γ-G_3n_/G_2n_** | **α-A_1n_/G-_1n_** | **γ-A_1n_/G_-1n_** | **α-G_L1_/A_L2_** | **γ-G_L1_/A_L2_** | **α-A_L2_/A_L3_** | **γ-A_L2_/A_L3_** |
| --- | --- | --- | --- | --- | --- | --- | --- | --- |
| *Free Kt-7* | | | | | | | | |
| **OL3(OPC)** | poor | poor | excellent | excellent | poor | poor | excellent | excellent |
| **OL3(SPC/E)** | poor | poor | excellent | excellent | poor | poor | excellent | excellent |
| **OL3(SPC/E)-1M** | poor | poor | excellent | excellent | poor | poor | excellent | excellent |
| **OL3_R2.7_** | poor | poor | good | good | poor | poor | excellent | excellent |
| **OL3_0BPh,CP_-gHBfix21** | poor | poor | good | good | poor | poor | excellent | excellent |
| **PAK** | poor | poor | excellent | excellent | poor | poor | excellent | excellent |
| **ROC** | poor | poor | excellent | excellent | poor | poor | excellent | excellent |
| **Chen&Garcia** | poor | poor | excellent | excellent | poor | poor | excellent | excellent |
| **DESRES** | poor | poor | good | good | poor | poor | good | excellent |
| **DES-Amber** | poor | poor | excellent | excellent | poor | poor | good | excellent |
| **BSSF1** | poor | poor | good | good | poor | poor | excellent | excellent |
| **CHARMM36** | poor | poor | poor | poor | poor | poor | poor | excellent |
| **CHARMM_Drude_** | poor | poor | poor | poor | poor | excellent | good | excellent |
| **AMOEBA** | poor | poor | excellent | excellent | excellent | excellent | excellent | excellent |
| *Kt-7 complexed with L7Ae* | | | | | | | | |
| **OL3(SPC/E)-L7Ae** | poor | poor | excellent | excellent | excellent | excellent | excellent | excellent |
| **AMOEBA-L7Ae** | poor | poor | excellent | excellent | excellent | excellent | excellent | excellent |

^a^The terms “excellent”, “good” and “poor” roughly summarize the observed simulation performance for the individual dihedrals.

Table S12: **Population analyses (in %) of the potassium binding-sites in L7Ae/Kt-7 protein-RNA complex simulations and their comparison with the free Kt-7.^a^**

| **OL3(SPC/E)** | | **OL3(SPC/E)-L7Ae** | |
| --- | --- | --- | --- |
| **Acceptor** | **Population** | **Acceptor** | **Population** |
| G_2n_(O6) | 150.3 | G_3n_(O6) | 149.9 |
| G_3n_(O6) | 145.8 | A_L3_(HO2') | 95.4 |
| G_3n_(N7) | 84.3 | A_L3_(O2') | 95.3 |
| G_4n_(N7) | 79.5 | G_2n_(O6) | 75.1 |
| G_4n_(OP2) | 67.4 | G_1b_(OP1) | 71.6 |
| G_5n_(O6) | 58.5 | G_4n_(N7) | 71.0 |
| G_4n_(O6) | 58.1 | A_L3_(O3') | 69.0 |
| G_1b_(N7) | 56.1 | G_3n_(N7) | 67.7 |
| G_2n_(N7) | 51.2 | G_5n_(O6) | 52.3 |
|  |  | G_4n_(O6) | 50.7 |
| **AMOEBA** | | **AMOEBA-L7Ae** | |
| **Acceptor** | **Population** | **Acceptor** | **Population** |
| G_3n_(O6) | 90.4 | G_3n_(O6) | 80.1 |
| G_2n_(O6) | 64.1 | G_3n_(N7) | 63.3 |
| G_4n_(OP2) | 50.0 | G_2n_(O6) | 62.2 |
| G_3n_(N7) | 50.0 | G_4n_(OP2) | 61.4 |

^a^Ion-binding sites with a population above 50% are listed. Populations of over 100% indicate more than one potassium ion being present at the site on average. Ion-binding sites appearing with both FFs are highlighted and color-coded for clarity.

Table S13: **Population analyses (in %) of the protein-RNA H-bonds formed in L7Ae/Kt-7 complex simulations.^a^**

| **OL3(SPC/E)** | | **AMOEBA** | |
| --- | --- | --- | --- |
| **H-bond (Donor - acceptor)** | **Population** | **H-bond (Donor - acceptor)** | **Population** |
| A_L3_(OP2) - Thr51(N) | 93.9 | Glu53(OE) - G_2n_(O2') | 87.1 |
| A_L3_(OP1) - Ala112(N) | 85.7 | A_L3_(OP2) - Thr51(OG1) | 81.9 |
| A_L3_(OP2) - Thr51(OG1) | 83.6 | G_3n_(OP2) - Lys56(NZ) | 71.6 |
| G_1b_(O6) - Glu53(N) | 67.5 | A_L3_(OP2) - Thr51(N) | 58.3 |
| Glu53(OE) - G_2n_(O2') | 55.7 | G_4n_(OP2) - Lys56(NZ) | 57.5 |
| G_3n_(OP2) - Lys56(NZ) | 43.2 | A_L3_(OP1) - Ala112(N) | 52.8 |
| Glu53(OE) - G_1b_(N2) | 38.7 | G_1b_(O6) - Glu53(N) | 50.7 |
| G_3n_(OP1) - Arg60(NH2) | 37.2 | A_L3_(N7) - Lys98(NZ) | 47.2 |
|  |  | Glu53(OE) - G_1b_(N1) | 33.6 |
| G_2n_(N7) - Lys56(NZ)^b^ | 19.7 | G_2n_(N7) - Lys56(NZ)^b^ | 5.4 |

^a^Protein-RNA H-bonds with population above 30% are listed.

^b^The G_2n_(N7) - Lys56(NZ) H-bond and generally the proximity of the lysine side-chain was suggested to cause a decrease in K^+^ occupancy of the major G_2n_(O6) binding site in OL3(SPC/E) FF (see the main text and Supporting Information S12). Because of its potential importance, the H-bond is listed in the Table even though it does not meet the population cut-off.

**Supporting Information Figures**

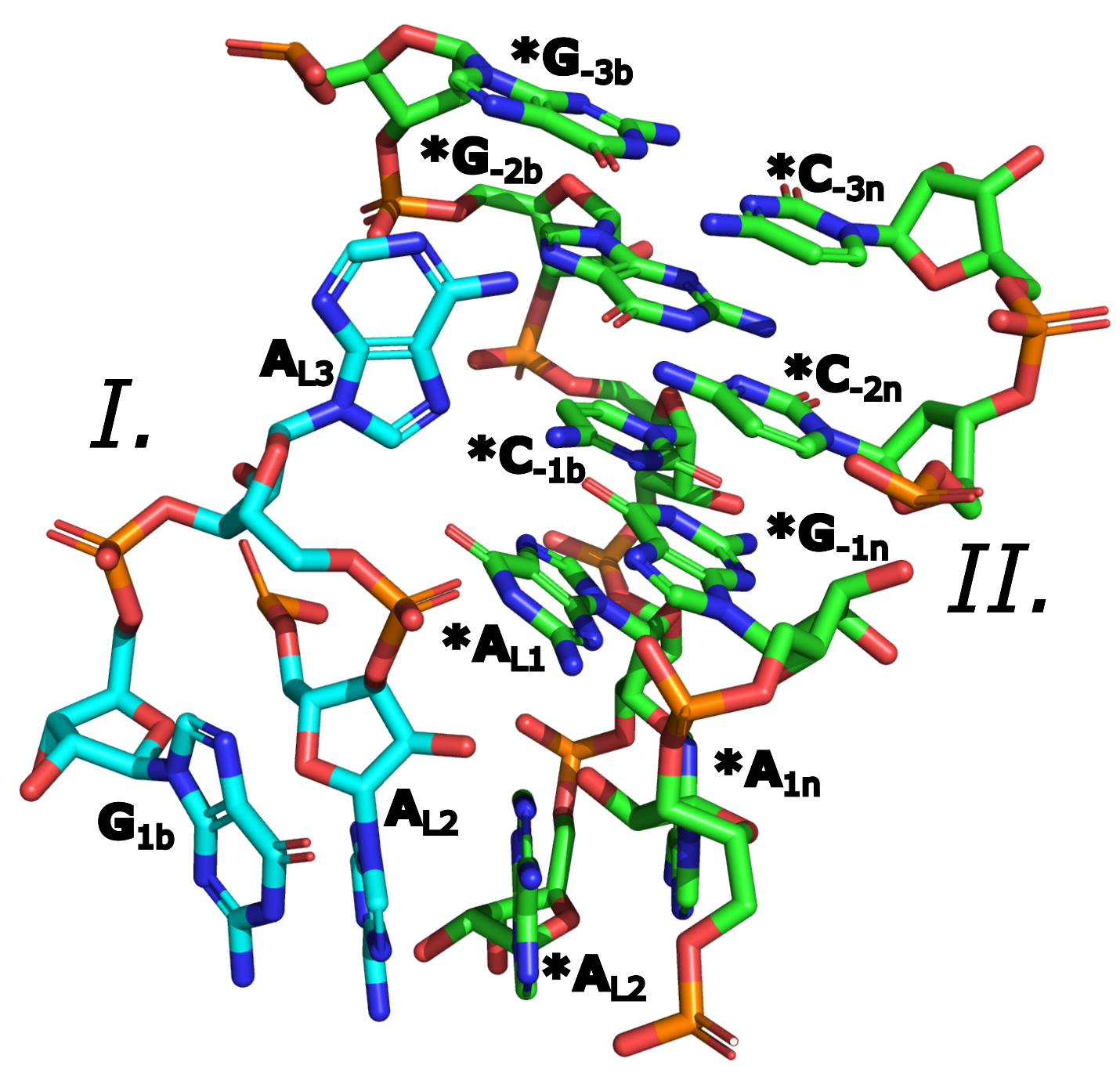

Figure S1: **Crystal packing contacts of A_L2_ in the X-ray structure of Kt-7.** Units I and II refer to the two parts of the kink-turn, related across the crystal lattice cell, with their carbon atoms colored cyan and green, respectively. Nucleotides in the crystallographic image are marked with an asterisk. The two A_L2_ bases from neighboring crystal lattice cells are stacking on top of each other. For clarity, only selected proximal residues are shown.

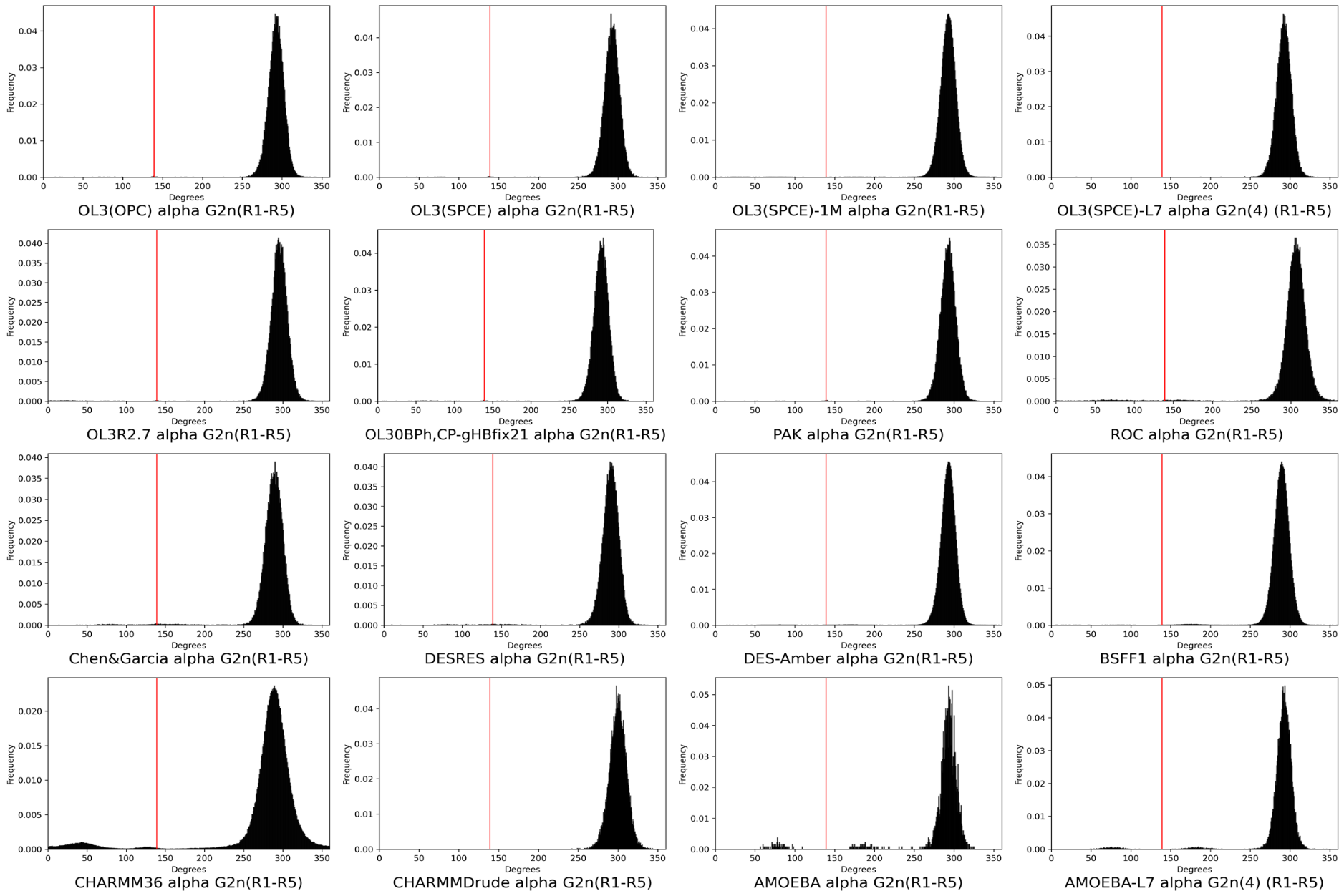

Figure S2A: **Histograms of the backbone dihedral α of suite G_3n_/G_2n_ for all tested FFs.** Values for combined simulation ensembles are shown. The vertical red line represents the experimental value (see Table S1).

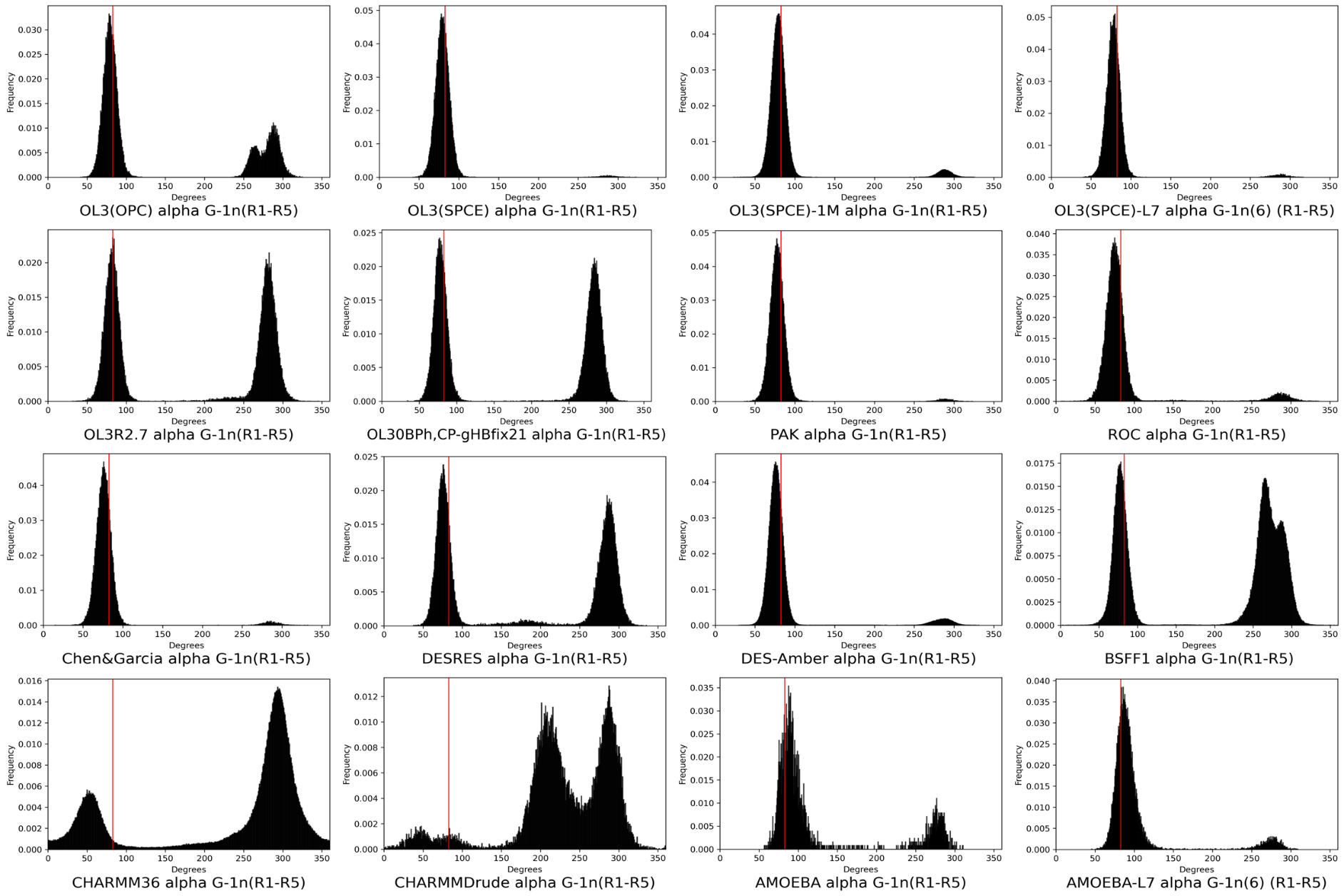

Figure S2B: **Histograms of the backbone dihedral α of suite A_1n_/G_-1n_ for all tested FFs.** Values for combined simulation ensembles are shown. The vertical red line represents the experimental value (see Table S1).

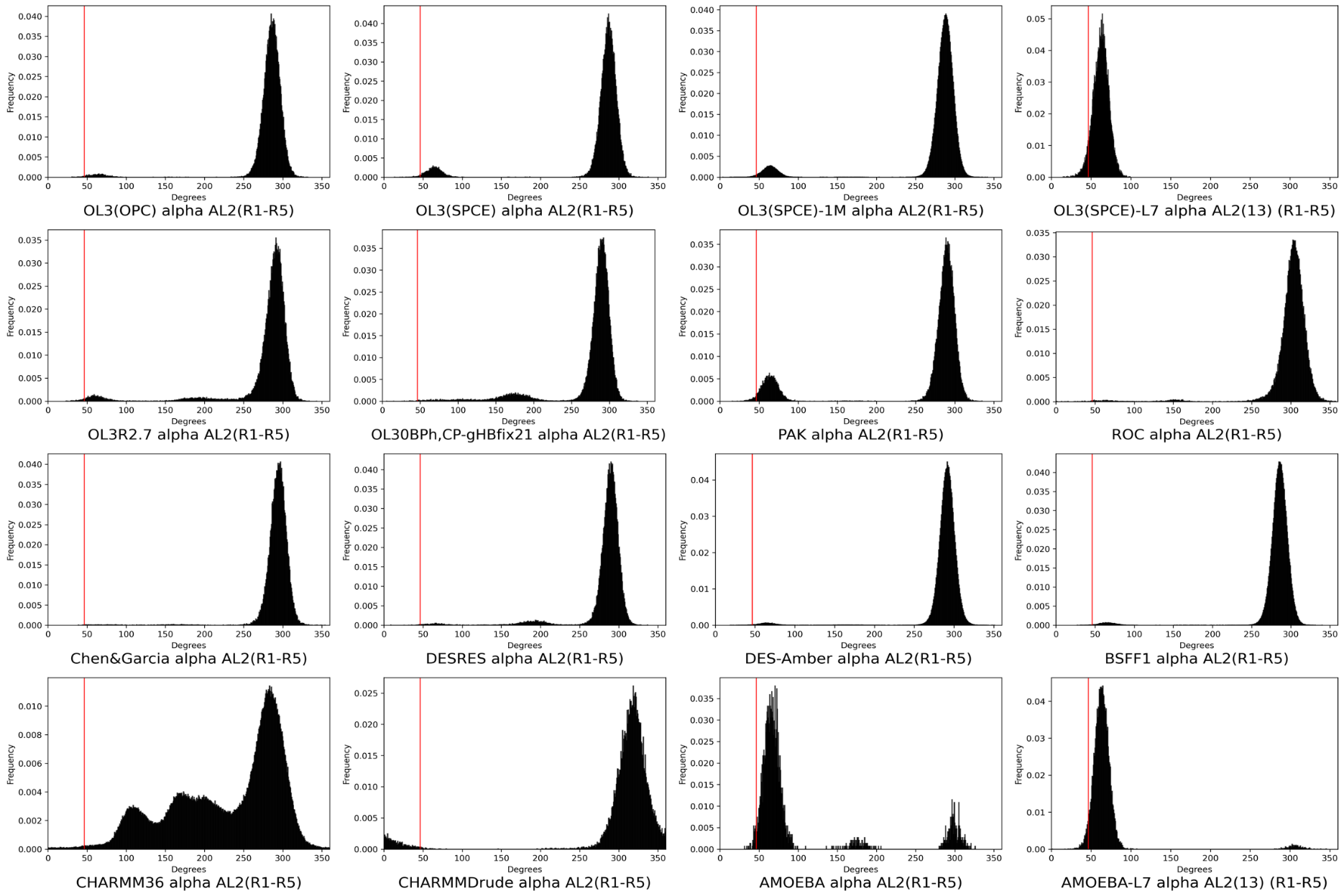

Figure S2C: **Histograms of the backbone dihedral α of suite G_L1_/A_L2_ for all tested FFs.** Values for combined simulation ensembles are shown. The vertical red line represents the experimental value (see Table S1)

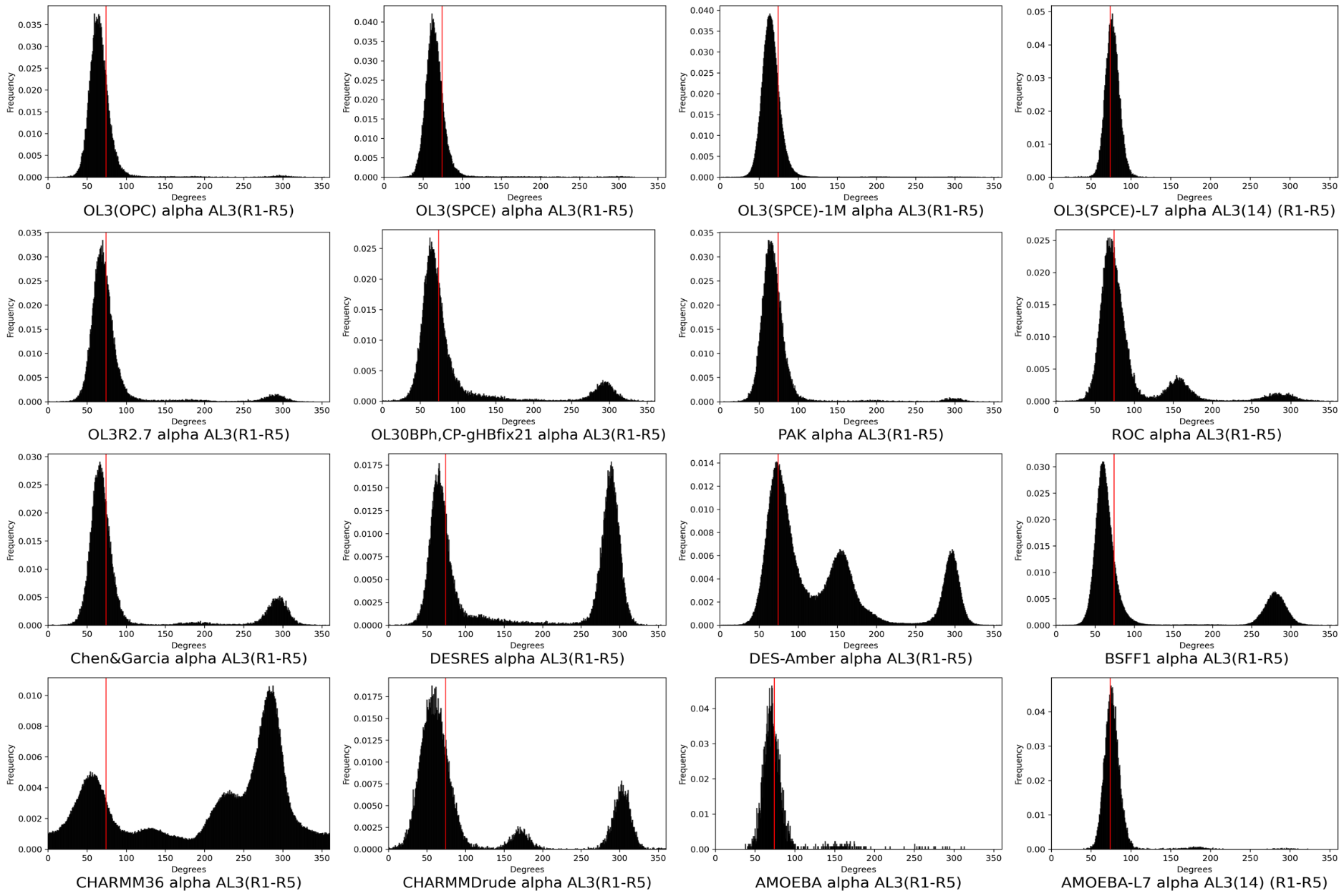

Figure S2D: **Histograms of the backbone dihedral α of suite A_L2_/A_L3_ for all tested FFs.** Values for combined simulation ensembles are shown. The vertical red line represents the experimental value (see Table S1).

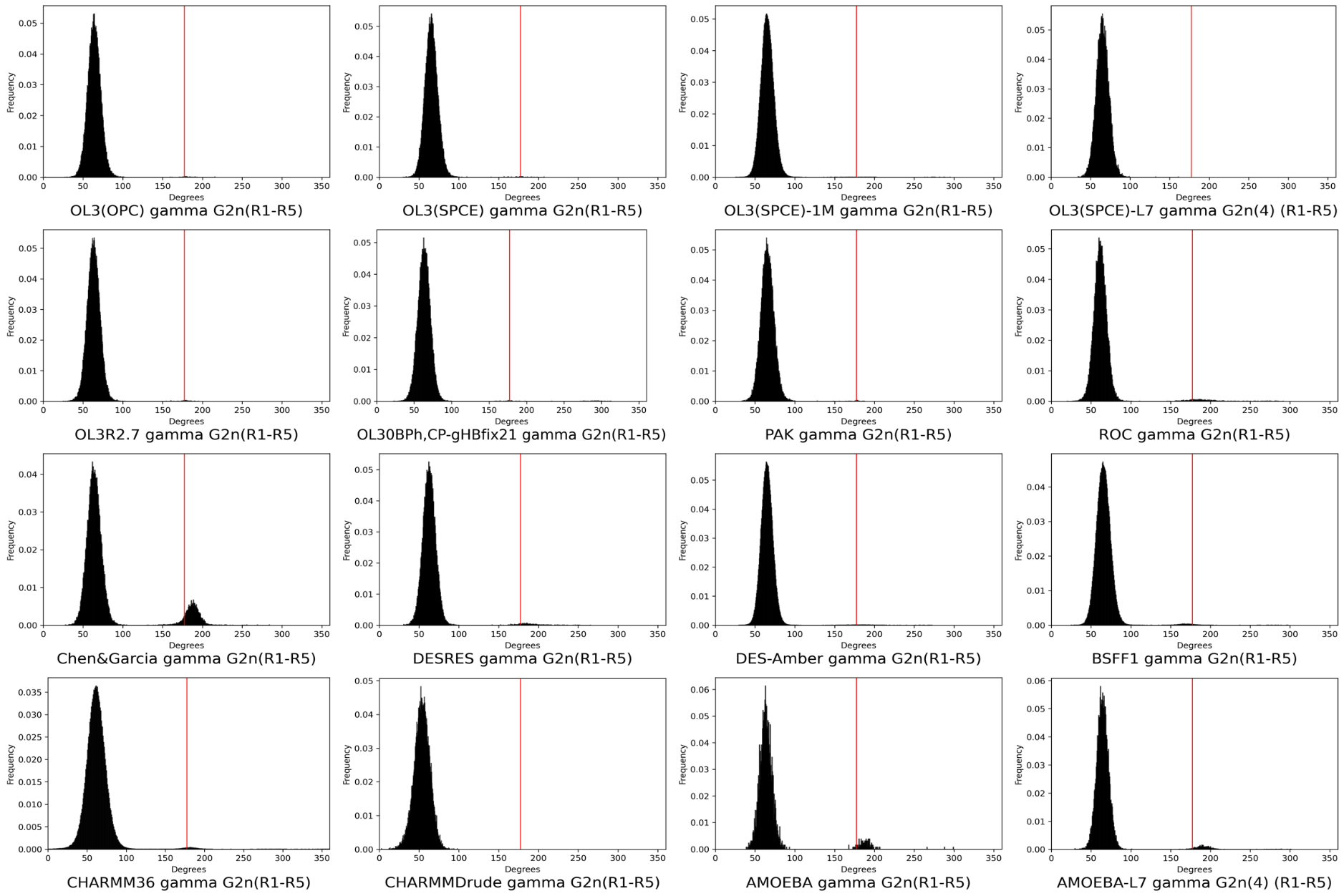

Figure S3A: **Histograms of the backbone dihedral γ of suite G_3n_/G_2n_ for all tested FFs.** Values for combined simulation ensembles are shown. The vertical red line represents the experimental value (see Table S1).

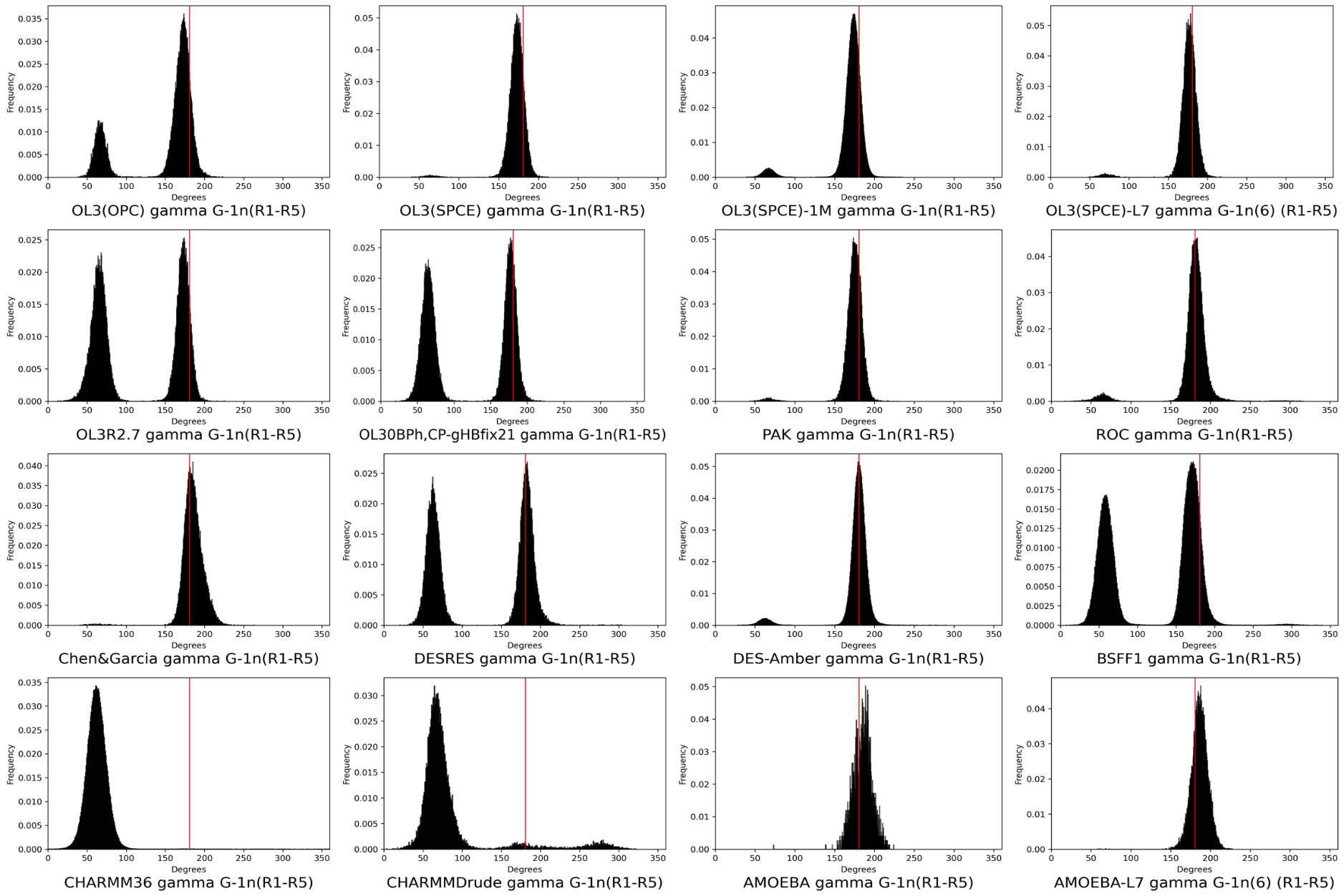

Figure S3B: **Histograms of the backbone dihedral γ of suite A_1n_/G_-1n_ for all tested FFs.** Values for combined simulation ensembles are shown. The vertical red line represents the experimental value (see Table S1).

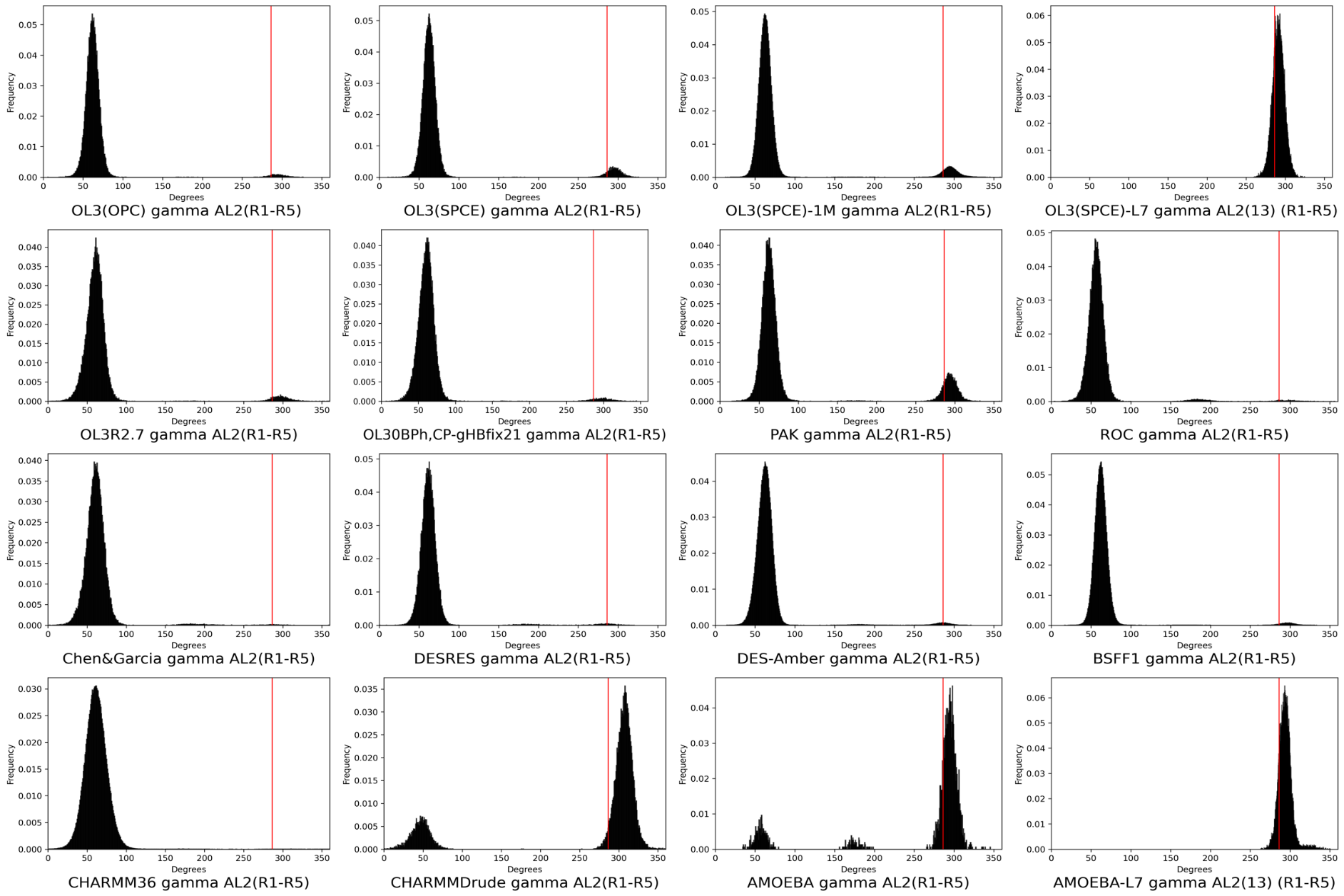

Figure S3C: **Histograms of the backbone dihedral γ of suite G_L1_/A_L2_ for all tested FFs.** Values for combined simulation ensembles are shown. The vertical red line represents the experimental value (see Table S1).

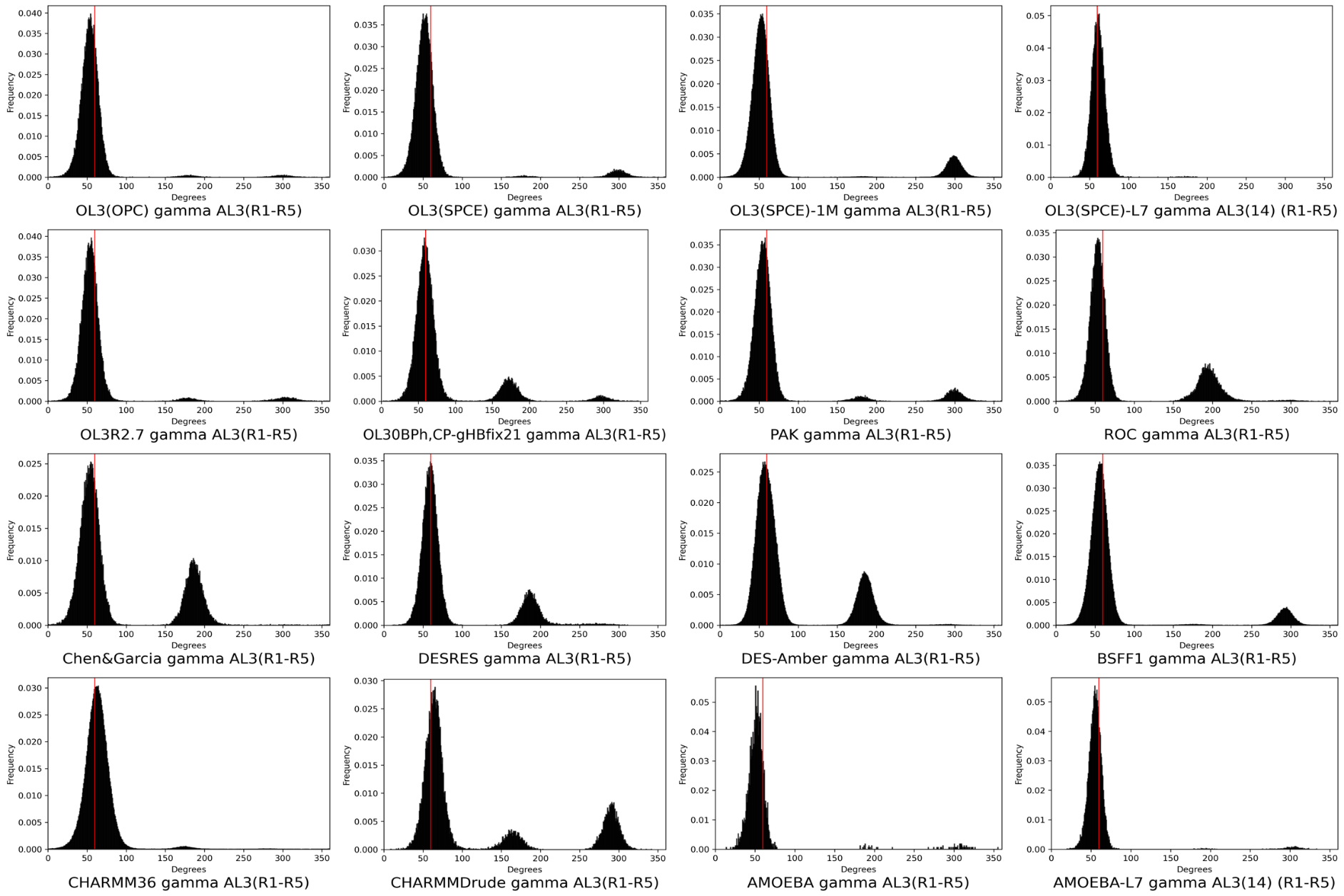

Figure S3D: **Histograms of the backbone dihedral γ of suite A_L2_/A_L3_ for all tested FFs.** Values for combined simulation ensembles are shown. The vertical red line represents the experimental value (see Table S1)

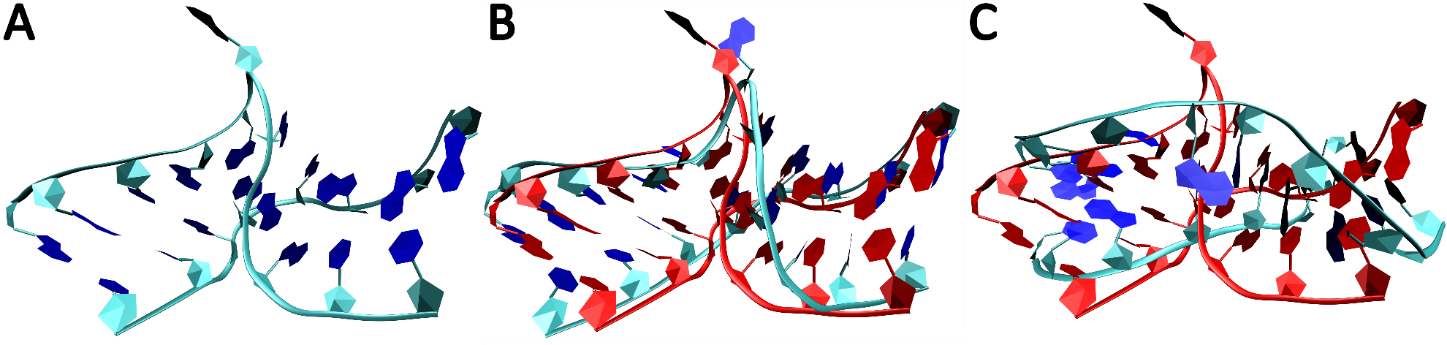

Figure S4: **Example of the Kt-7 unkinking.** (A) The starting structure; (B) Slightly straightened intermediary conformation with the stems still intact (7.37 µs); (C) Unkinked structure with progressive deterioration of the stems (9.68 µs). All simulation figures are taken from a replicate 4 simulation with the DESRES FF. For panels B and C, the starting structure is also shown as a red overlay.

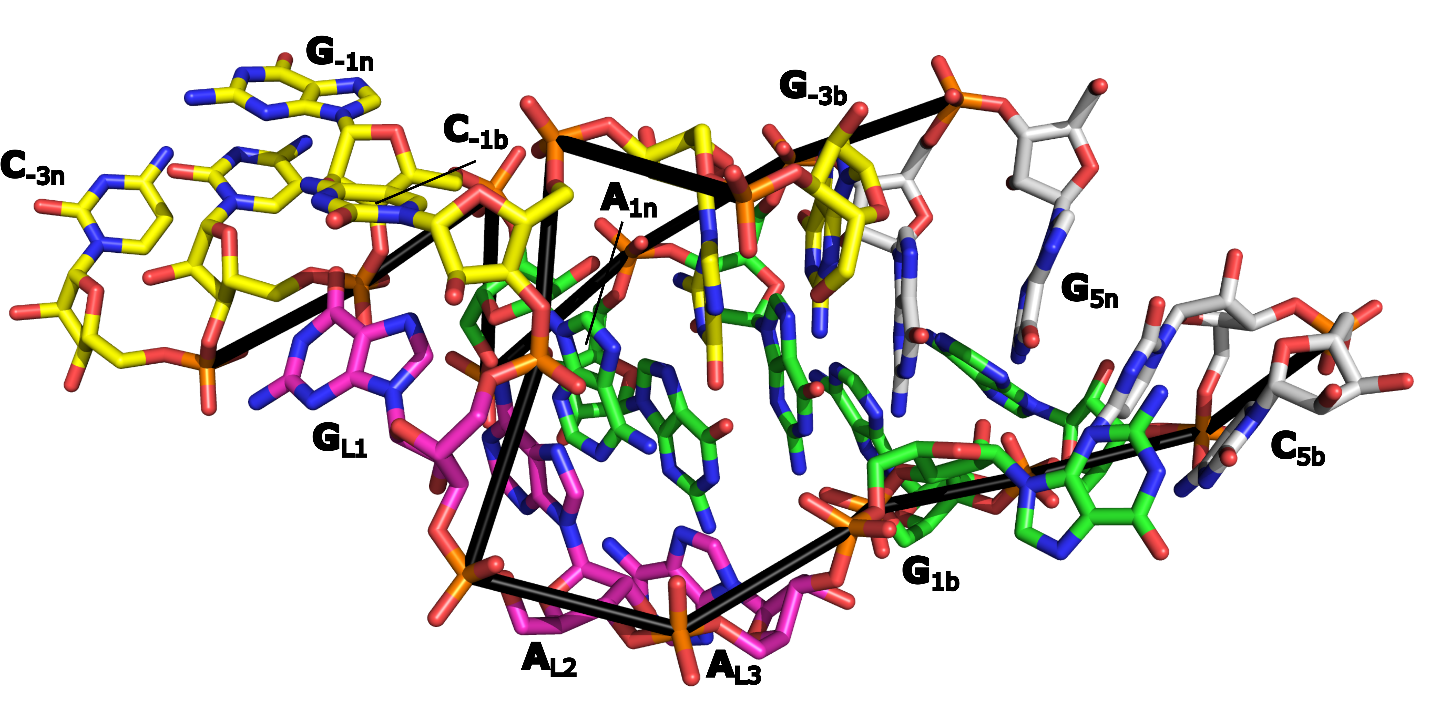

Figure S5: **Severely** **disrupted Kt-7 structure observed in CHARMM36 simulations.** Virtually all the native H-bonds of both stems are all lost. The structure shown is from replicate 4 (3.03 µs).

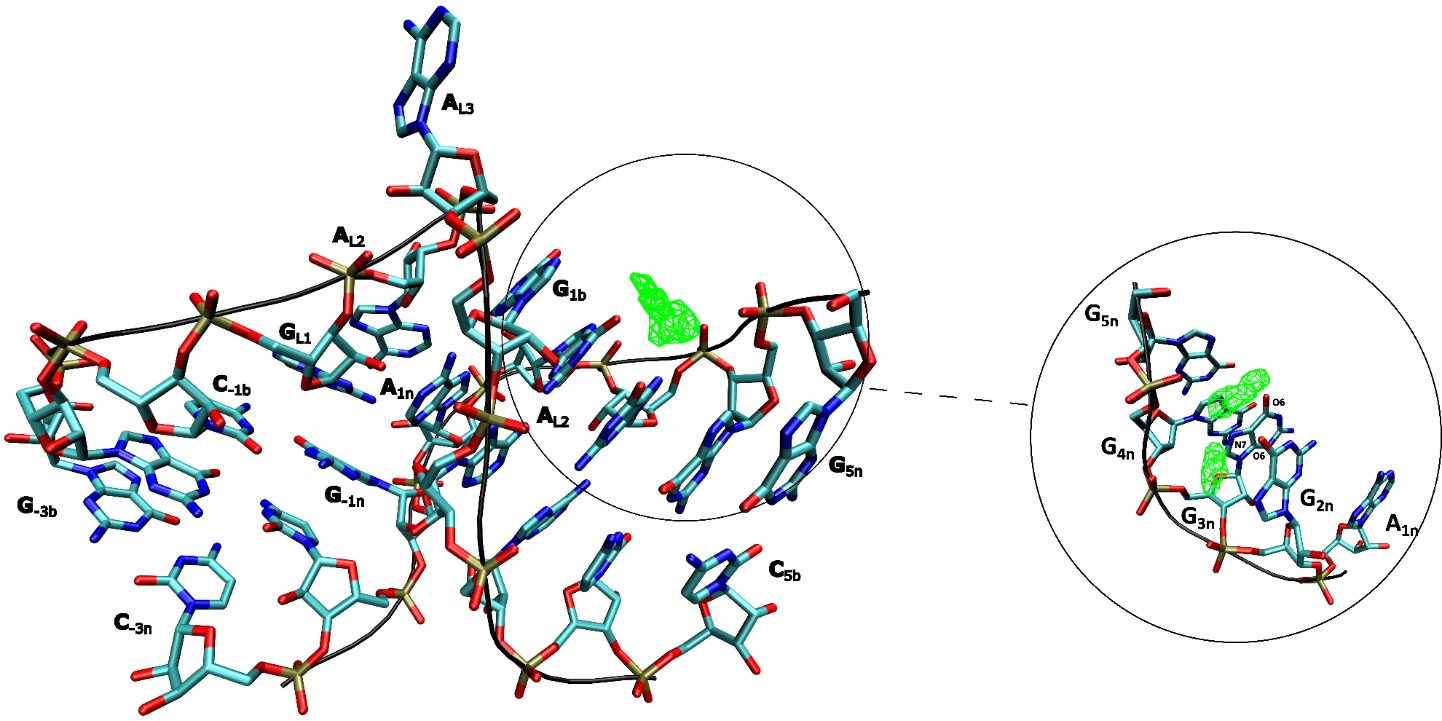

Figure S6: **Visualization of the most densely occupied cation binding sites in MD simulations of Kt-7**. The density map of K^+^ ions (shown as green wireframe) reveals the ion-binding sites near atoms G_2n_(O6), G_3n_(O6) and G_3n_(N7) (shown in the inset). Carbon atoms are colored in cyan. The shown density map was generated based on one of the OL3(OPC) simulations, however, similar positioning of the ion binding sites was observed with all FFs albeit their exact populations could differ.

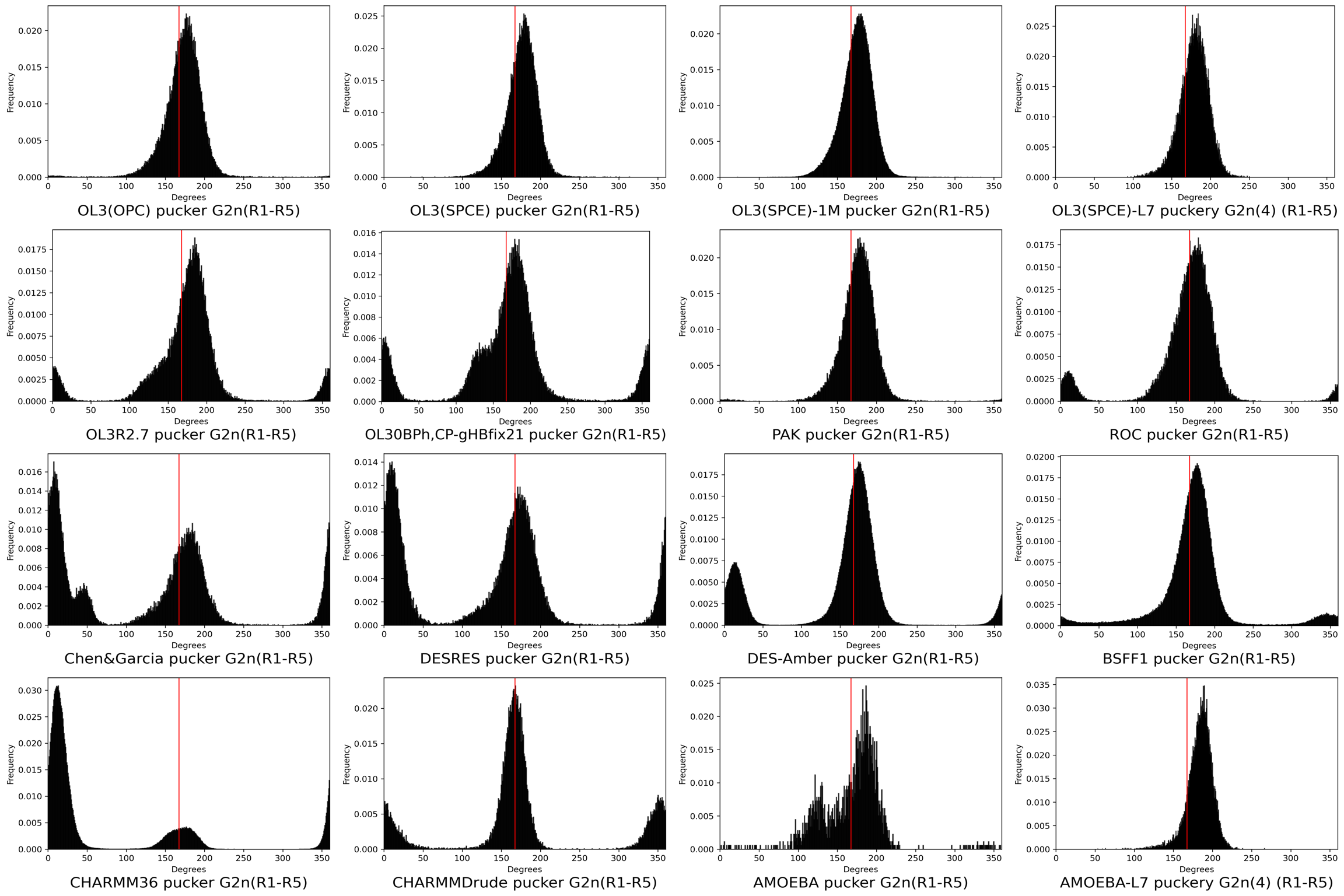

Figure S7A: **Histograms of the pucker of G_2n_ for all tested FFs.** Values for combined simulation ensembles are shown. The vertical red line represents the experimental value (see Table S1).

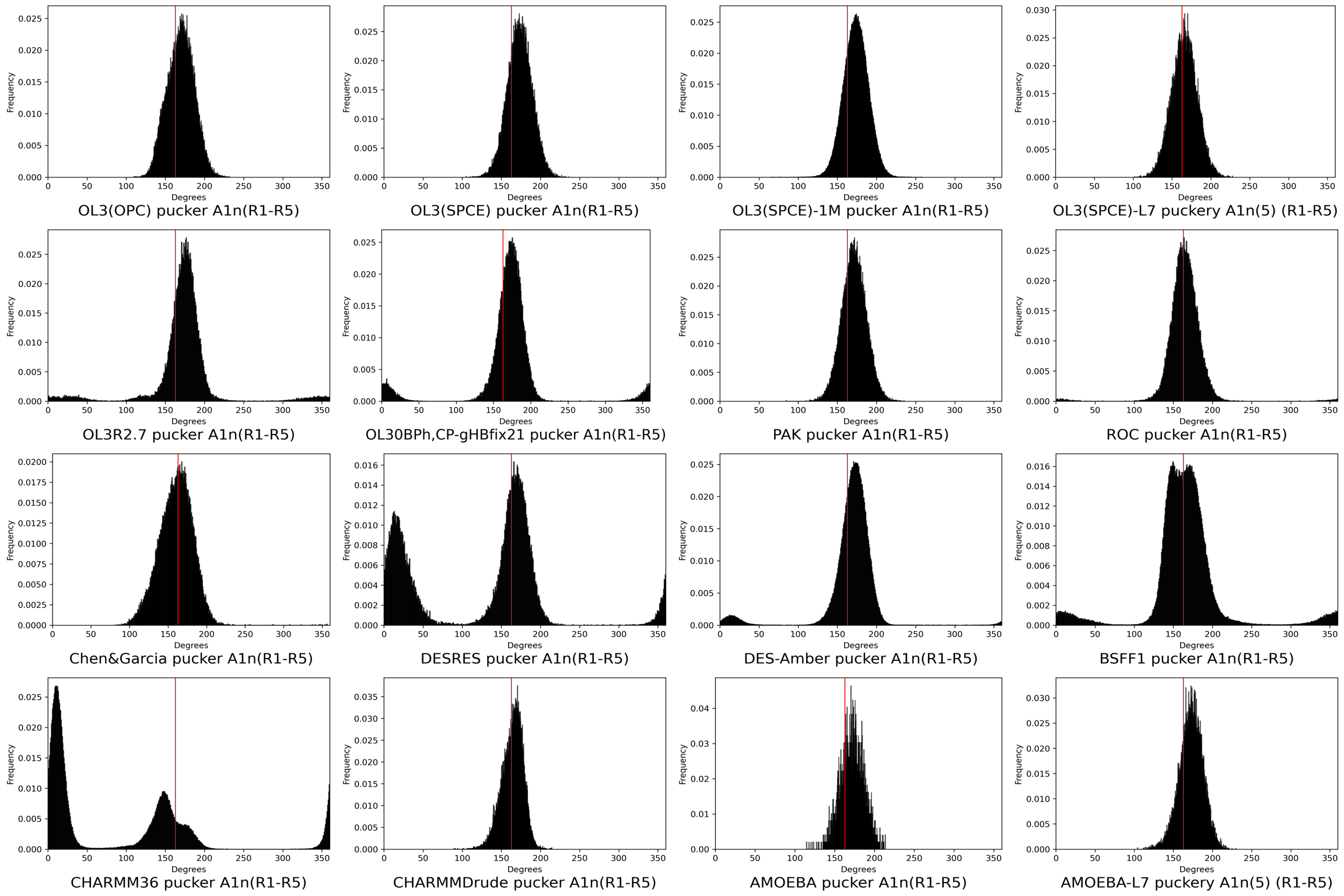

Figure S7B: **Histograms of the pucker of A_1n_ for all tested FFs.** Values for combined simulation ensembles are shown. The vertical red line represents the experimental value (see Table S1).

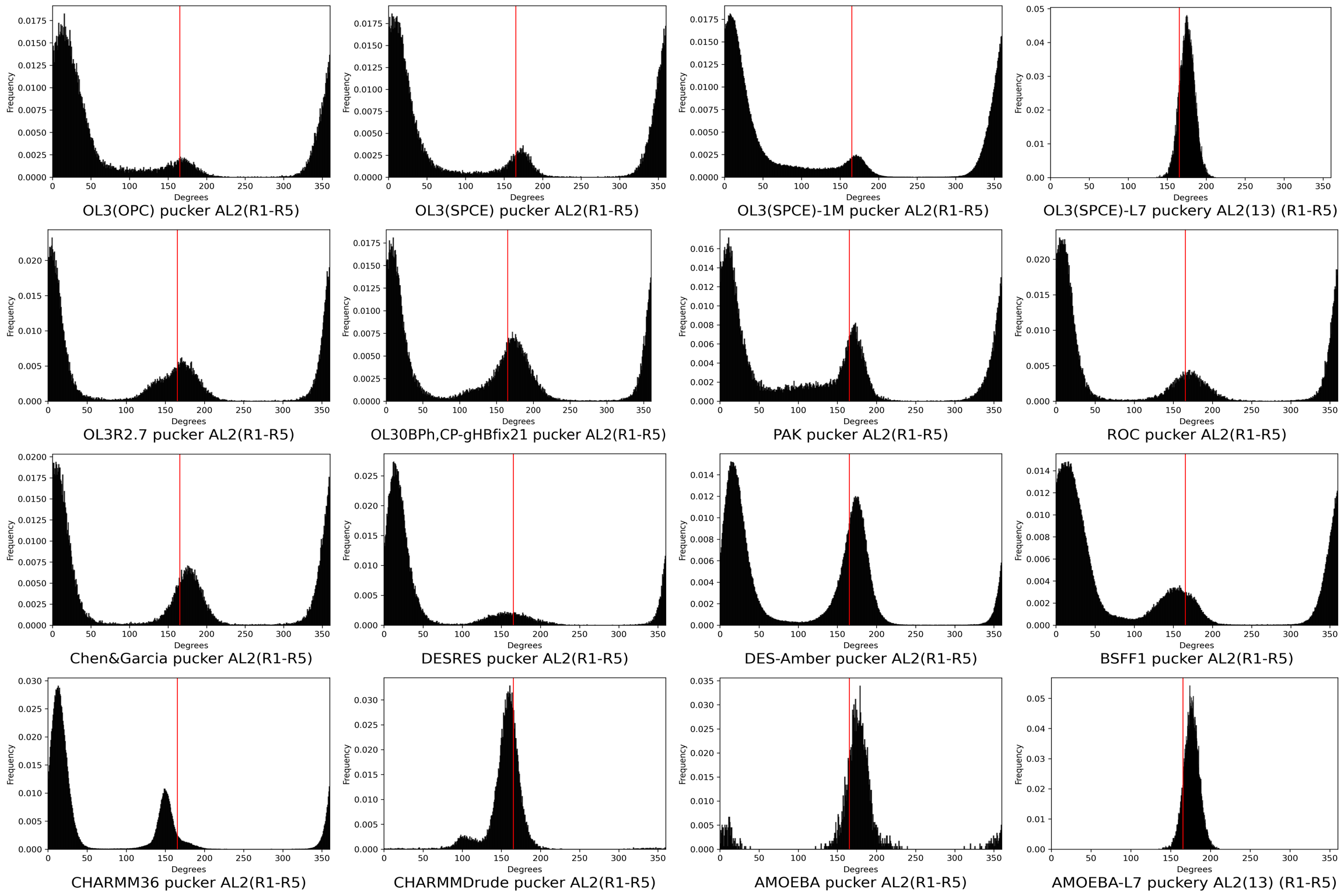

Figure S7C: **Histograms of the pucker of A_L2_ for all tested FFs.** Values for combined simulation ensembles are shown. The vertical red line represents the experimental value (see Table S1).

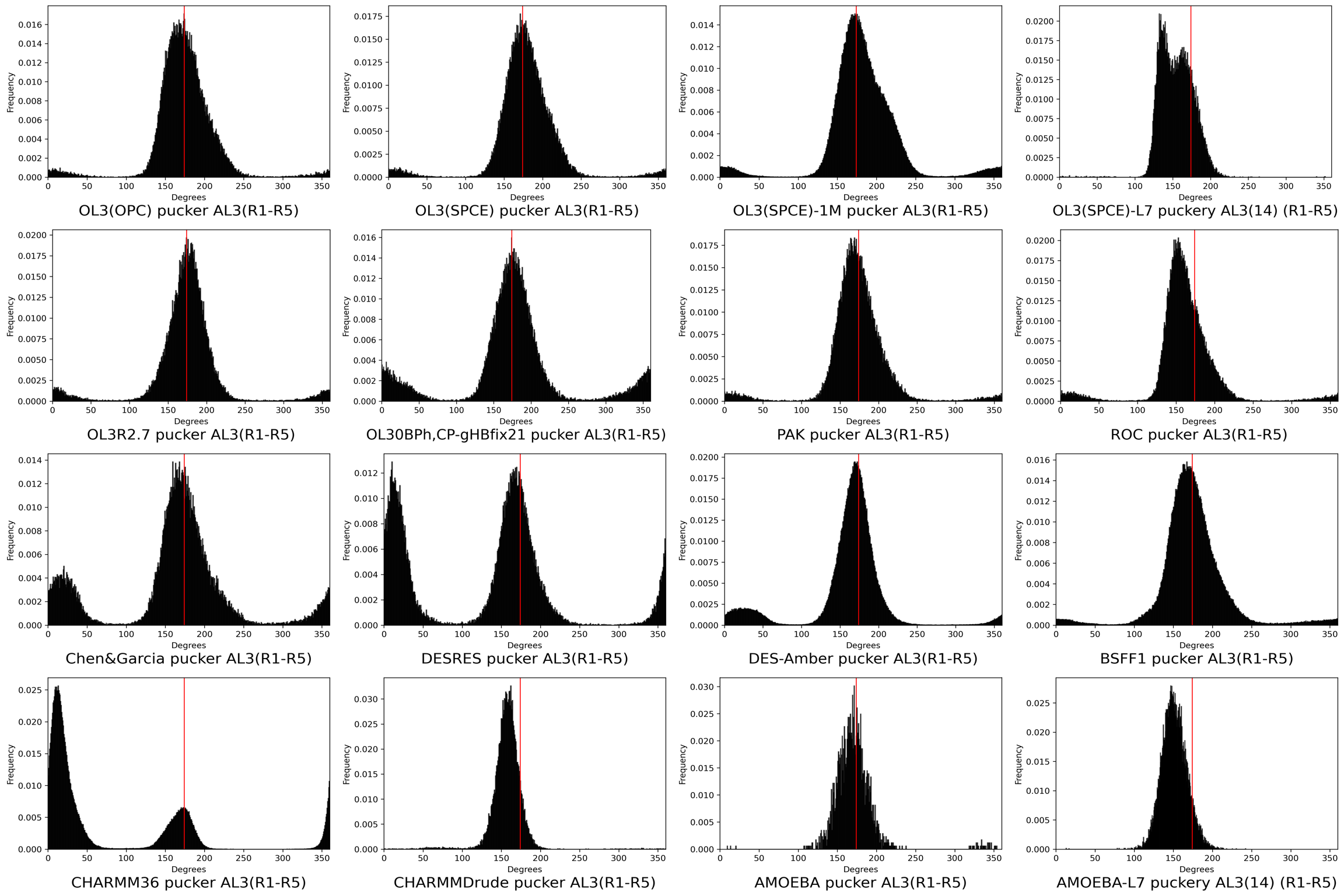

Figure S7D: **Histograms of the pucker of A_L3_ for all tested FFs.** Values for combined simulation ensembles are shown. The vertical red line represents the experimental value (see Table S1).

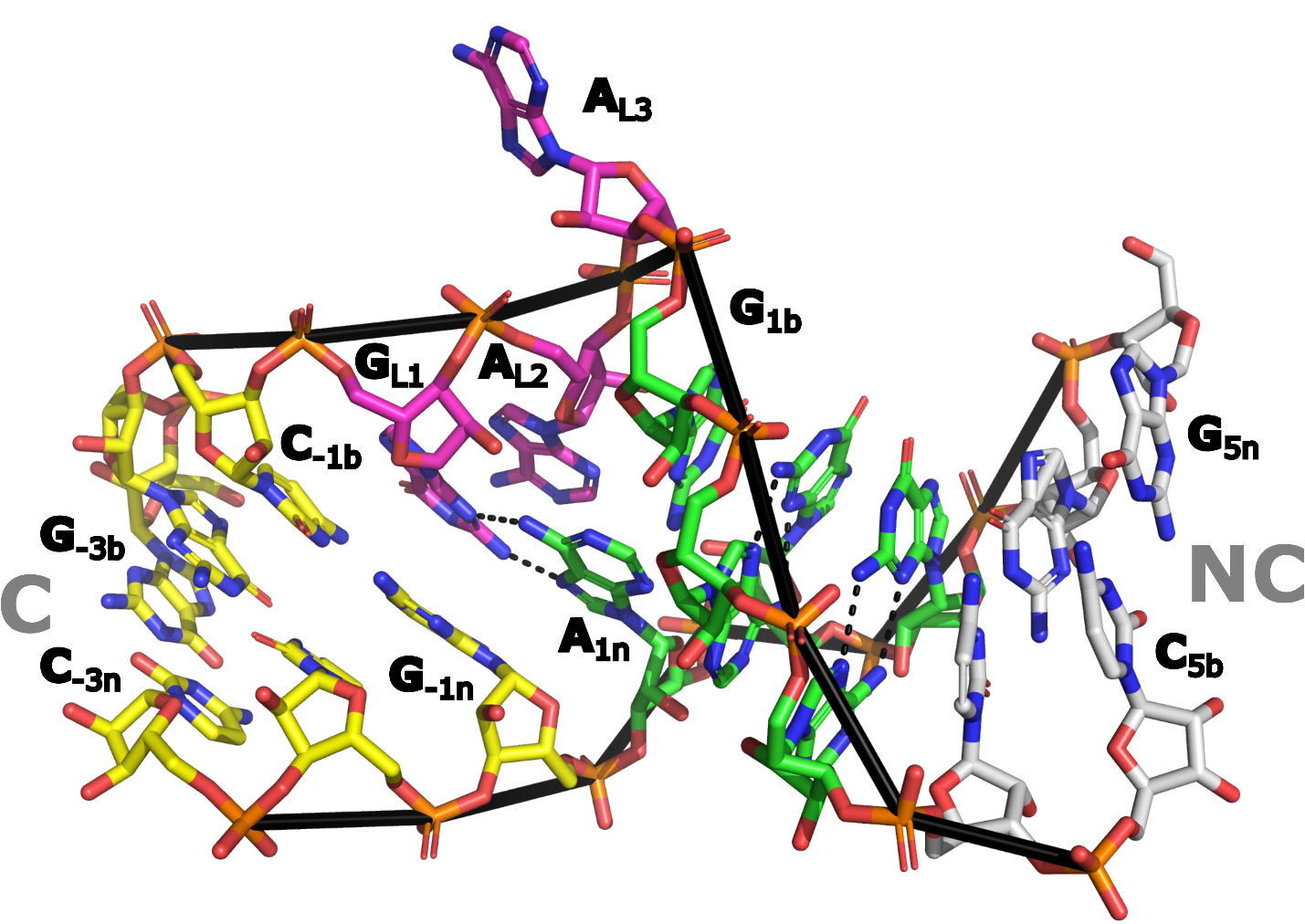

Figure S8: **Formation of a spurious, non‑native AG base pair between A_1n_ and G_L1_.** In several force fields (OL3_0BPh, CP‑_gHBfix21, DESRES, and DES‑Amber), the native G_1b_–A_1n_ base pair breaks during unkinking, subsequently leading to formation of a spurious AG base pair between A_1n_ and G_L1_. The snapshot shown was taken from the DESRES simulation, replicate 1 (1.26 µs), with residues labeled as in Figure 1 (main text) and the AG base pair H-bonds shown with dashed black lines.

### **Supporting Information References**

(1) Huang, L.; Lilley, D. M. The kink turn, a key architectural element in RNA structure. *J. Mol. Biol.* **2016**, *428* (5), 790-801. DOI: <https://doi.org/10.1016/j.jmb.2015.09.026>.

(2) Falb, M.; Amata, I.; Gabel, F.; Simon, B.; Carlomagno, T. Structure of the K-turn U4 RNA: a combined NMR and SANS study. *Nucleic Acids Res.* **2010**, *38* (18), 6274-6285. DOI: 10.1093/nar/gkq380 (acccessed 3/31/2025).

(3) Altona, C.; Sundaralingam, M. Conformational analysis of the sugar ring in nucleosides and nucleotides. New description using the concept of pseudorotation. *J. Am. Chem. Soc.* **1972**, *94* (23), 8205-8212. DOI: 10.1021/ja00778a043.
